# Live imaging and biophysical modeling support a button-based mechanism of somatic homolog pairing in *Drosophila*

**DOI:** 10.1101/2020.08.30.265108

**Authors:** Myron Child, Jack R. Bateman, Amir Jahangiri, Armando Reimer, Nicholas C. Lammers, Nica Sabouni, Diego Villamarin, Grace C. McKenzie-Smith, Justine E. Johnson, Daniel Jost, Hernan G. Garcia

## Abstract

The spatial configuration of the eukaryotic genome is organized and dynamic, providing the structural basis for regulated gene expression in living cells. In *Drosophila melanogaster*, 3D genome organization is characterized by somatic homolog pairing, where homologous chromosomes are intimately paired from end to end; however, the process by which homologs identify one another and pair has remained mysterious. A recent model proposed that specifically interacting “buttons” encoded along the lengths of homologous chromosomes drive somatic homolog pairing. Here, we turned this hypothesis into a precise biophysical model to demonstrate that a button-based mechanism can lead to chromosome-wide pairing. We tested our model and constrained its free parameters using live-imaging measurements of chromosomal loci tagged with the MS2 and PP7 nascent RNA labeling systems. Our analysis showed strong agreement between model predictions and experiments in the separation dynamics of tagged homologous loci as they transition from unpaired to paired states, and in the percentage of nuclei that become paired as embryonic development proceeds. In sum, as a result of this dialogue between theory and experiment, our data strongly support a button-based mechanism of somatic homolog pairing in *Drosophila* and provide a theoretical framework for revealing the molecular identity and regulation of buttons.

## Introduction

Eukaryotic genomes are highly organized within the three-dimensional volume of the nucleus, from the large scale of chromosome territories to the smaller-scale patterned folding of chromosomal segments called Topologically Associated Domains (TADs) and the association of active and inactive chromatin into separate compartments (Szabo et al., 2019). Disruption of these organizational structures can have large consequences for gene expression and genome stability (Despang et al., 2019; Kragesteen et al., 2018; Lupiáñez et al., 2015; Rosin et al., 2019), emphasizing the importance of fully understanding the mechanisms underlying three-dimensional genome organization.

While many principles of genome organization are common among eukaryotes, differences have been noted between organisms and cell types. For example, in somatic cells in *Drosophila*, an additional layer of nuclear organization exists: homologous chromosomes are closely juxtaposed from end to end, a phenomenon known as somatic homolog pairing (Joyce et al., 2016; Stevens, 1908). While similar interchromosomal interactions occur transiently in somatic cells of other species and during early meiotic phases of most sexually reproducing eukaryotes, the widespread and stable pairing of homologous chromosomes in somatic cells of *Drosophila* appears to be unique to Dipteran flies (Joyce et al., 2016; King et al., 2019; McKee, 2004). Notably, the close juxtaposition of paired homologs can have a dramatic impact on gene expression through a process known as transvection, whereby regulatory elements on one chromosome influence chromatin and gene expression on a paired chromosome (Duncan, 2002; Fukaya and Levine, 2017). Although somatic homolog pairing was first described over 100 years ago (Stevens, 1908), the molecular mechanisms by which homologous chromosomes identify one another and pair have yet to be described.

During the early stages of *Drosophila* development, maternal and paternal genomes are initially separated and become paired as embryogenesis proceeds. Prior analyses of the initiation of somatic homolog pairing have relied primarily on DNA fluorescent *in situ* hybridization (DNA-FISH) to label homologous loci in fixed embryos, and have led to a model in which somatic homolog pairing slowly increases with developmental time through independent associations along the lengths of each chromosome arm (Figure 1A) (Fung et al., 1998; Gemkow et al., 1998; Hiraoka et al., 1993). This model is further supported by recent studies that converged on a “button” model for pairing, which hypothesizes that pairing is initiated at discrete sites along the length of each chromosome (Figure 1B) (Rowley et al., 2019; Viets et al., 2019). However, the molecular nature of these hypothesized buttons is as yet unclear, nor is it clear whether this proposed model could lead to *de novo* pairing in the absence of some unknown active process that identifies and pairs homologous loci.

**Figure 1:**
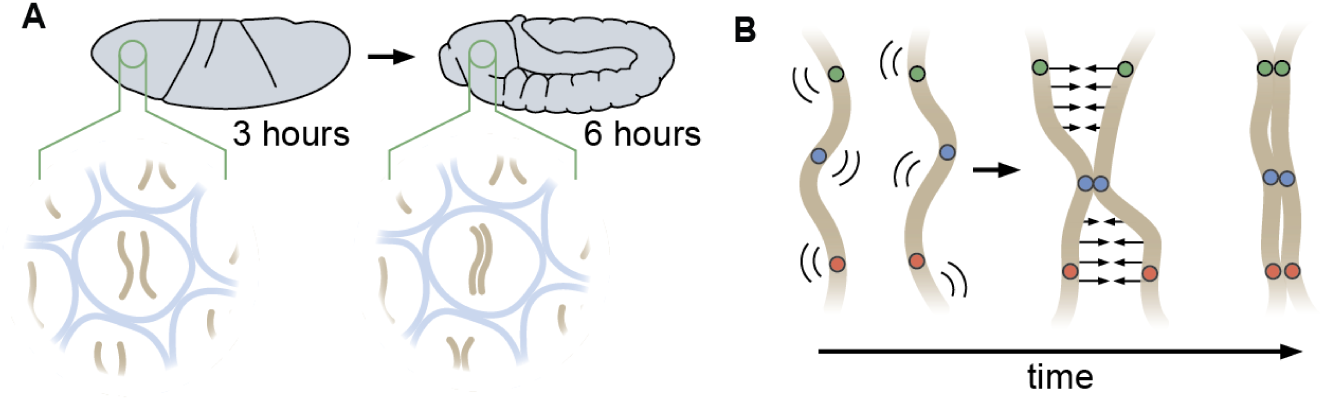
Schematic of homologous chromosome pairing in somatic cells in *D. melanogaster*. (A) Over the course of embryonic development, homologous chromosomes pair along their lengths. (B) Button model for homolog pairing in which each chromosome carries a series of sites that have affinity for the same site on a homologous chromosome.

Here we turned the “button” mechanism for somatic homolog pairing into a precise biophysical model that defines parameters for the activities of pairing buttons, informed by observations of pairing dynamics in living cells. Our simulations showed that chromosome-wide pairing can be established through random encounters between specifically interacting buttons that are dispersed across homologous chromosomes at various possible densities using a range of binding energies that are reasonable for protein-protein interactions. Importantly, we found that active processes are not necessary to explain pairing via our model, as all of the interactions necessary for stable pairing are initiated by reversible random encounters that are propagated chromosome-wide. We tested our model and constrained its free parameters by assessing its ability to predict pairing dynamics measured via live imaging. Our model successfully predicted that, once paired, homologous loci remain together in a highly stable state. Further, the model also accurately predicted the dynamics of pairing through the early development of the embryo, as measured by the percentage of nuclei that become paired as development proceeds and by the dynamic interaction of individual loci as they transition from unpaired to paired states. In sum, through an interplay between theory and experiment aimed at probing molecular mechanisms, our analysis provides quantitative data that strongly support a button model as the underlying mechanism of somatic homolog pairing and establishes the conceptual infrastructure to uncover the molecular identity, functional underpinnings, and regulation of these buttons.

## Results

### 1) FORMALIZING A BUTTON-BASED POLYMER MODEL OF HOMOLOGOUS PAIRING

Prior studies have suggested that somatic homolog pairing in *Drosophila* may operate via a button mechanism between homologous loci (AlHaj Abed et al., 2019; Erceg et al., 2019; Fung et al., 1998; Gemkow et al., 1998; Rowley et al., 2019; Viets et al., 2019). In this model, discrete regions capable of pairing specifically with their corresponding homologous segments are interspersed throughout the chromosome. To quantitatively assess the feasibility of a button mechanism, we implemented a biophysical model of homologous pairing (Fig. 2A). Briefly, we modeled homologous chromosome arms as polymers whose dynamics are driven by short-range, attractive, specific interactions between homologous loci (buttons) to account for pairing (Materials and Methods). These buttons are present at a density *ϱ* along the chromosome and bind specifically to each other with an energy *E*_*p*_. We included short-range, non-specific interactions among (peri)centromeric regions to account for large-scale HP1-mediated clustering of centromeres (Materials and Methods), which may also impact global genome organization inside nuclei (Rosin et al., 2018; Strom et al., 2017) and thus may affect pairing. As initial conditions for our simulations, we generated chromosome configurations with all centromeres at one pole of the nucleus (a ‘Rabl’ configuration; Supplementary Movies 1 and 2), typical of early embryonic fly nuclei (Dernburg et al., 1996). To account for the potential steric hindrance of non-homologous chromosomes that could impede pairing, we simulated two pairs of homologous polymers.

**Figure 2:**
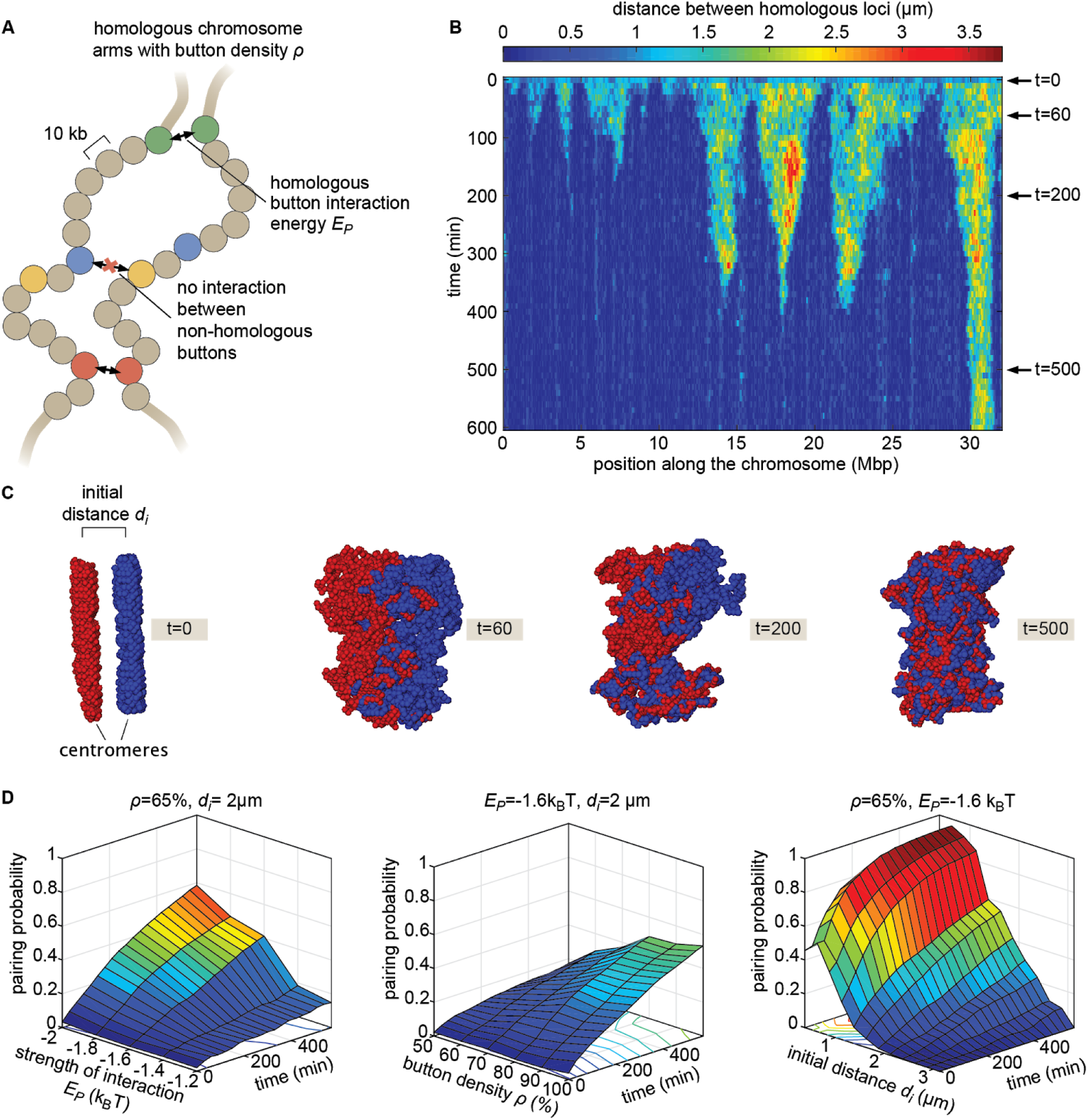
The homologous button model. (A) Pairing between homologous chromosomes is assumed to be driven by specific, short-range attractive interactions of strength *E*_*p*_ between certain homologous regions, named buttons. Each 10-kb monomer in the simulation corresponds to one locus. (B) Kymograph of the time-evolution of the distances between homologous regions predicted by the model in one representative simulated stochastic trajectory for a button density of *ϱ=*65%, an interaction strength of *E*_*p*_ =−1.6k_B_T, and an initial distance *d*_*i*_ = 1 μ*m*. See Figure 2 – Supplementary Figure 1 for other examples for various *d*_*i*_ values. (C) Snapshots of the pair of homologous chromosomes at various time points along the simulation in (B) (see also Supplementary Movies 1 and 2). (D) Predicted average pairing probability between euchromatic homologous loci (considered as paired if their distance ≤ 1 μ*m*) as a function of time and of the strength of interaction *E*_*p*_ (left), the button density *ϱ* (center), and the initial distance *d*_*i*_ between homologous chromosomes (right).

When we monitored the distances between homologous loci in our simulations as a function of time (Fig. 2B and Fig. 2 – Supp. Fig. 1), qualitatively we observed that this thermodynamic model can lead to the time-progressive pairing of homologous chromosomes (Fig. 2B) and the gradual intermingling of the two homologous chromosome territories (Fig. 2C). Pairing in our simulations operates via a stochastic zipping process: once random fluctuations lead to the pairing of one pair of homologous loci, the pairing of nearest-neighbor buttons is facilitated along the lengths of the homologous chromosomes in a zipper-like manner (Fig. 2B). Full chromosome-wide pairing results from the progression of many zippers that “fire” at random positions and times along the chromosome.

We systematically investigated the roles of button density along the genome *ϱ*, of the strength of the pairing interaction *E*_*p*_, and of the initial distance between homologous chromosomes *d*_*i*_ in dictating pairing dynamics (Fig. 2D). For a given density, there is a critical value of *E*_*p*_ below which no large-scale pairing event occurs independently of the initial conditions (Fig. 2 – Sup. Fig. 2A) since pairing imposes a huge entropic cost for the polymers and thus requires a sufficient amount of energy to be stabilized. Beyond this critical point, higher strengths of interactions and higher button densities lead to faster and stronger pairing (Fig. 2D left, center). We also find that the non-specific interactions among (peri)centromeric regions included in our model facilitate pairing, but that such interactions are not strictly necessary (Fig. 2 – Sup. Fig. 2D).

The initial spatial organization of chromosomes also strongly impacts pairing efficiency. When homologous chromosomes are initially far apart, pairing is dramatically slowed and impaired (Fig. 2D, right) due to the presence of the other simulated chromosomes between them (Fig. 2 – Sup. Fig. 2B). We also observed that our initial chromosome configurations corresponding to a Rabl-like organization (with all centromeres at one pole of the nucleus) promotes pairing by allowing homologous buttons to start roughly aligned (Marshall and Fung, 2016; Nicodemi et al., 2008; Penfold et al., 2012) (Fig. 2 – Sup. Fig. 2F). Taken together, these systematic analyses of model parameters support the view that the homologous button model is compatible with pairing. These results are fully consistent with a similar button-like polymer model developed to investigate pairing during yeast meiosis (Marshall and Fung, 2016).

As an alternative model, we asked whether buttons that interact non-specifically could also explain somatic pairing. We simulated the dynamics of polymers having such non-specific buttons and never observed significant chromosome-wide pairing (Fig. 2 – Sup. Fig. 2C,E). These results are complementary to previous works in which we showed that the weak, non-specific interactions between epigenomic domains that drive TAD and compartment formation in *Drosophila* (Ghosh and Jost, 2018; Jost et al., 2014) cannot establish and maintain stable pairing by themselves (Pal et al., 2019). Thus, in addition to button density, interaction strength, and initial organization of chromosomes, a key mechanism for pairing is the specificity of preferential interactions between homologous regions.

### 2) LIVE IMAGING REVEALS HOMOLOGOUS PAIRING DYNAMICS

The button model in Figure 2 makes precise predictions about pairing dynamics at single loci along the chromosome. To inform the parameters of the model and to test its predictions, it is necessary to measure pairing dynamics in real time at individual loci of a living embryo. To do so, we employed the MS2/MCP (Bertrand et al., 1998) and PP7/PCP (Chao et al., 2008) systems for labeling nascent transcripts. Here, each locus contains MS2 or PP7 loops that can be visualized with distinct colors in living embryos (Chen et al., 2018; Fukaya et al., 2016; Garcia et al., 2013; Lim et al., 2018). Specifically, we designed transgenes encoding MS2 or PP7 loops under the control of UAS (Brand and Perrimon, 1993) and integrated them at equivalent positions on homologous chromosomes (Fig. 3A). Activation of transcription with GAL4 creates nascent transcripts encoding the MS2 or PP7 stem loops, each of which can be directly visualized by maternally providing fluorescently labeled MCP (MCP-mCherry) or PCP (PCP-GFP) in the embryo. The accumulation of fluorescent molecules on nascent transcripts was detected via laser-scanning confocal microscopy, providing relative three-dimensional positions of actively transcribing chromosomal loci in living *Drosophila* embryos (Chen et al., 2018; Lim et al., 2018).

**Figure 3:**
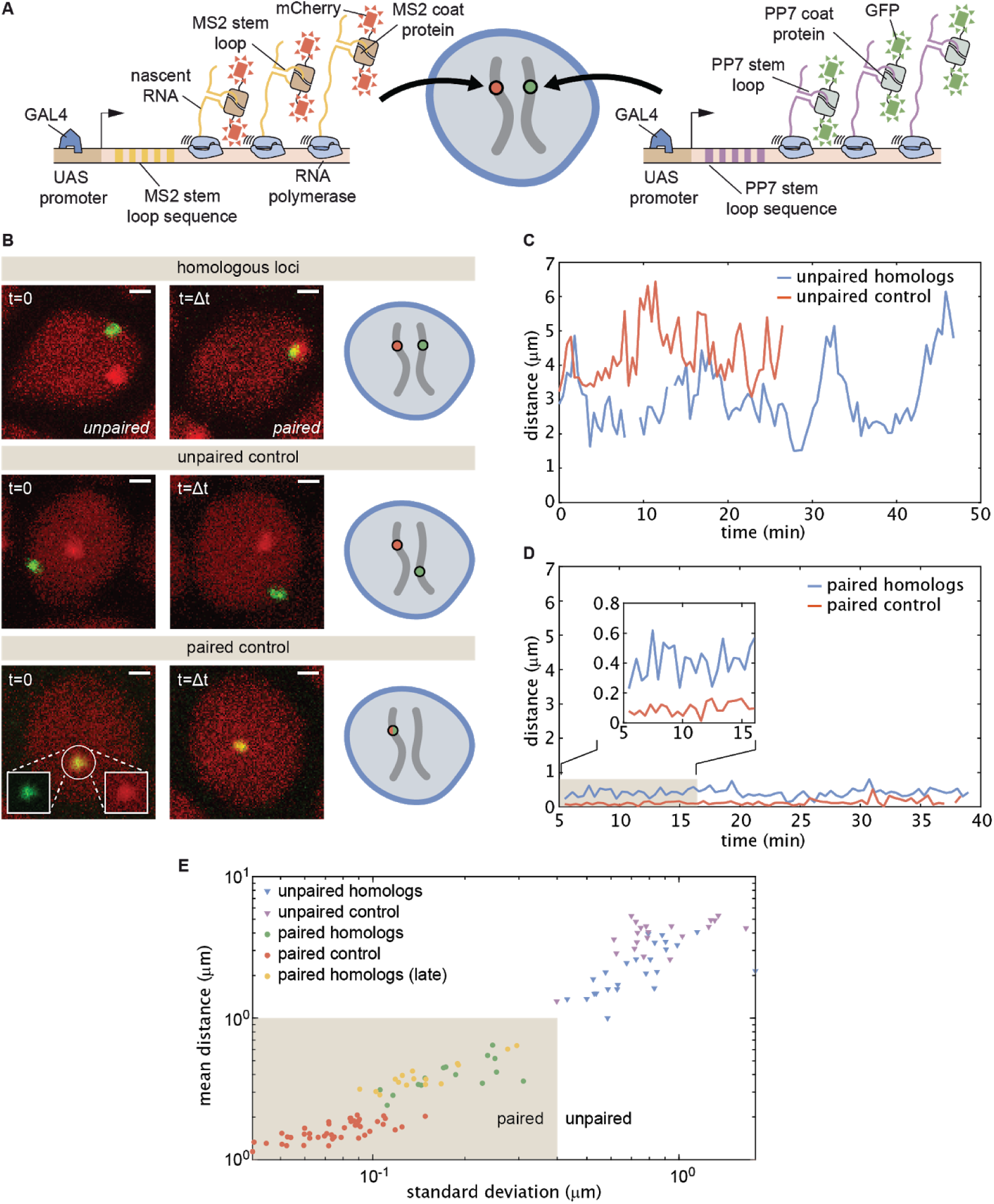
Live imaging of chromosomal loci provides dynamic single-locus spatiotemporal information about somatic homolog pairing. (A) Schematic of the MS2 and PP7 nascent mRNA labeling scheme for live imaging of homologous loci. Expression of the stem loops is driven by UAS under the control of a GAL4 driver. (B) Snapshots at two time points from homologous chromosomal loci with one allele tagged with MS2 and one allele tagged with PP7 (top), negative control consisting of non-homologous loci labeled with MS2 and PP7 (middle), and positive controls correspond to a single reporter containing interlaced MS2 and PP7 stem loops on the same chromosome (bottom). Scale bars represent 1 µm. See also Supplementary Movies 3-6. (C) Representative traces of the dynamics of the distance between imaged loci for unpaired homologous loci and the negative control showing how both loci pairs have comparable distance dynamics. (D) Representative traces of the dynamics of the distance between imaged loci for paired homologous loci and the positive control demonstrating how the distance between paired loci is systematically higher than the control. (E) Mean and SD of the distance between reporter transgenes, where each data point represents a measurement over the length of time that the loci were imaged (10-50 min). The shaded region indicates the criterion used to define whether homologs are paired (< 1.0 µm mean distance, < 0.4 µm SD) based on the distribution of points where homologs were qualitatively assessed as paired (yellow and green points).

We focused on embryos that had completed the maternal-to-zygotic transition and began to undergo gastrulation at approximately 2.5-5 h after embryo fertilization, when pairing begins to increase substantially (Fung et al., 1998). We integrated transgenes into two genomic locations on chromosome 2 at polytene positions 38F and 53F, and analyzed embryos with MS2 and PP7 loops at the same positions on homologous chromosomes in order to monitor pairing (Fig. 3B, top; Supplementary Movies 3 and 4; Materials and Methods). As a negative control, we imaged embryos in which loops were integrated at various positions on homologous chromosomes (MS2 at position 38F and PP7 at position 53F), where we expect no pairing between transgenes (Fig. 3B, middle; Supplementary Movie 5). Finally, as a positive control for the spatial colocalization of MS2 and PP7 loops, we analyzed embryos where MS2 and PP7 loops were interlaced in a single transgene (Chen et al., 2018; Wu et al., 2014) on one chromosome at polytene position 38F (Fig. 3B, bottom; Supplementary Movie 6). For each case, we imaged multiple embryos for 30-60 min, and used custom MATLAB scripts to determine the relative 3D distances between chromosomal loci over time (Materials and Methods).

In embryos with both PP7 and MS2 transgenes integrated at polytene position 38F (Fig. 3B, top), the majority of nuclei could be qualitatively classified into one of two categories. In “unpaired” nuclei, homologous loci were typically separated by >1 µm with large and rapid changes in inter-homolog distances (e.g. Fig. 3C, blue), with a mean distance of 2.2 µm and standard deviation (SD) of 1.2 µm averaged over 30 nuclei. The measured mean distance between homologous loci was comparable within error, though systematically smaller, than the mean distance between loci in the negative control, where transgenes were integrated at nonhomologous positions (Fig. 3C, red, mean distance = 4.0 µm, SD = 1.3 µm, n = 21 nuclei). In contrast, in “paired” nuclei, homologous loci remained consistently close to one another over time, with smaller dynamic changes in inter-homolog distance (Fig. 3D, blue, mean distance = 0.4 µm, SD = 0.3 µm, n = 25 nuclei). Interestingly, while the diffraction-limited signals produced from homologous loci occasionally overlapped in paired nuclei, their average separation was systematically larger than that of the positive-control embryos carrying interlaced MS2 and PP7 loops (Fig. 3D, red, mean distance = 0.2 µm, SD = 0.1 µm, n = 44 nuclei). This control measurement also constitutes a baseline for the experimental error of our quantification of inter-homolog distances (Chen et al., 2018). Our measurements thus confirmed previous observations of transgene pairing in the early embryo in which signals from paired loci maintained close association but did not completely coincide over time (Lim et al., 2018). Notably, of 38 nuclei qualitatively scored as having paired homologs, we never observed a transition back to the unpaired state over a combined imaging time of more than 8 h. Embryos with PP7 and MS2 transgenes integrated in homologous chromosomes at polytene position 53F showed comparable dynamics of inter-homolog distances for nuclei in unpaired and paired states (Fig. 3 – Sup. Fig. 1A, B). Thus, somatic homolog pairing is a highly stable state characterized by small dynamic changes in the distance between homologous loci.

Our assessment thus far has been based on a qualitative definition of pairing. In order to devise a stringent quantitative definition of homologous pairing, we measured inter-transgene distances for homologous loci as well as for the unpaired and paired controls throughout gastrulation. We also included measurements from older embryos (∼11-12 h after fertilization) using the driver *R38A04-GAL4* (Jenett et al. 2012) to express the transgenes in epidermal cells, where pairing is expected to be widespread (Fung et al., 1998; Gemkow et al., 1998). We measured the mean and SD of the inter-transgene distance for each nucleus over ∼10-50 min. From these data, we established a quantitative and dynamic definition of somatic homolog pairing based on a mean distance <1.0 µm and a corresponding standard deviation <0.4 µm (Fig. 3E, shaded region). By this definition, we considered paired 100% of nuclei that we had qualitatively scored as such, but excluded all nuclei scored as unpaired. As expected, this definition also scored 100% (15/15) of the tracked nuclei from older embryos as paired. Data for paired nuclei from early versus late embryos were in close agreement (Fig. 3E, yellow), demonstrating that pairing observed in early embryos is representative of pairing during later stages of development.

We next analyzed the progression of pairing through the first 6 h of development in single embryos carrying MS2 and PP7 transgenes in homologous chromosomes at positions 38F and 53F. To accomplish this goal, we collected data for short (∼10 min) intervals every 30 min from 2.5 h to 6 h of development, and analyzed inter-homolog distances as outlined above. We then plotted the mean of this distance as a function of its SD for each nucleus analyzed at each time point to create a dynamic assessment of somatic homolog pairing over developmental time. As expected, we detected an overall decrease in mean inter-homolog distance and its SD as development progressed (Fig. 4A, Fig. 3 – Sup. Fig. 1C). To directly compare our analysis to prior studies, we binned nuclei into paired and unpaired states based on their mean and SD as defined in Figure 3E and plotted the percentage of paired nuclei at each developmental time point (Fig. 4B). Consistent with previous literature (Fung et al., 1998; Gemkow et al., 1998), we observed a steady increase in the proportion of paired nuclei (Fig. 4B); however, by our dynamical definition of pairing, the percentage of nuclei that are paired is systematically lower at most timepoints than results using DNA-FISH (Fig. 4 – Sup. Fig. 1). This disagreement likely reflects differences between the classic, static definition of pairing based on overlapping DNA-FISH signals in the one frame accessible by fixed-tissue measurements as opposed to our dynamics-based definition, which demands that loci be paired over several consecutive frames. In sum, we have demonstrated that our system captures the progression of somatic homolog pairing over developmental time, making it possible to contrast theoretical predictions and experimental measurements.

**Figure 4.**
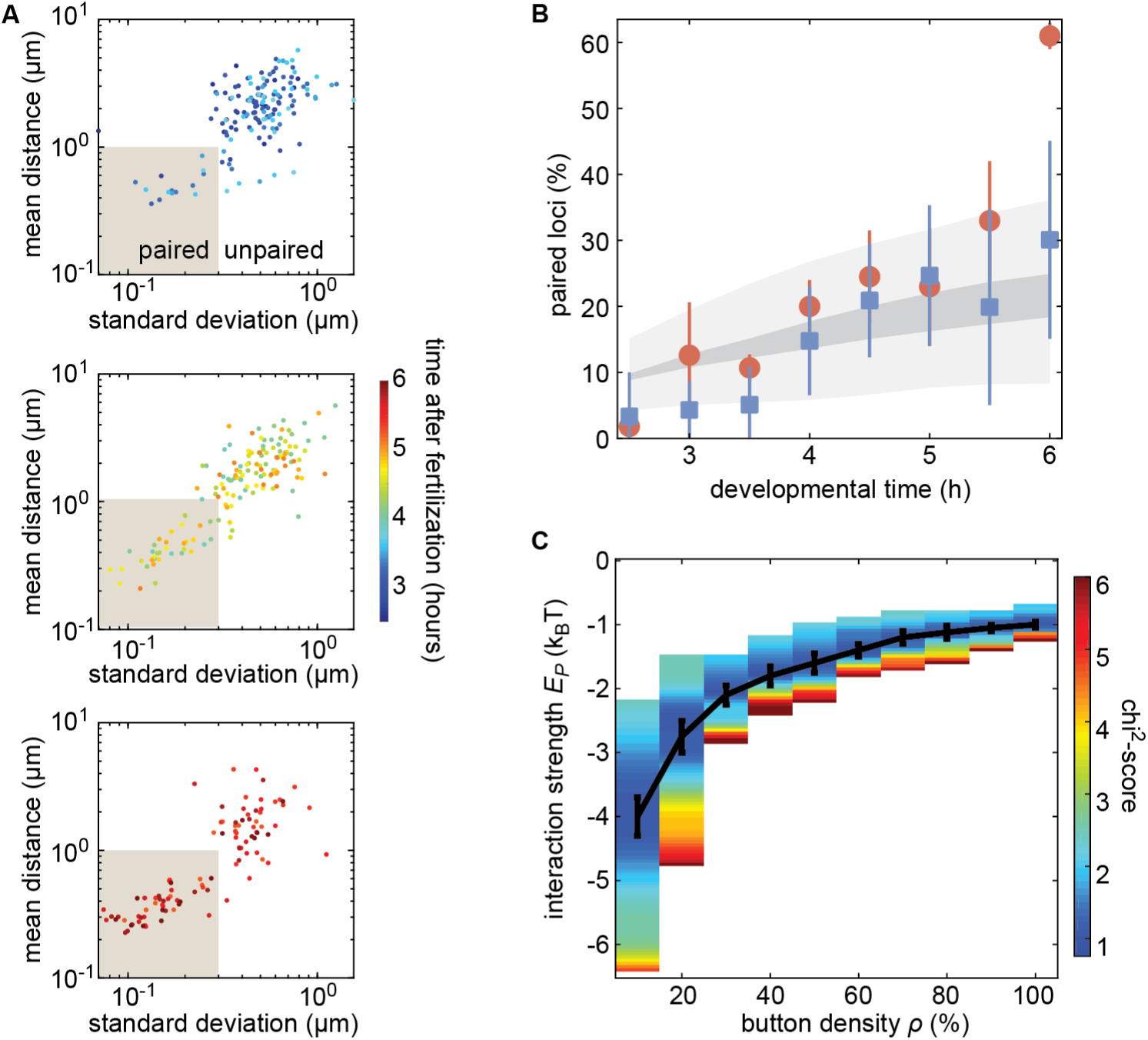
The homologous button model recapitulates the observed developmental dynamics of pairing. (A) Mean and SD of the separation of each pair of transgenes integrated at position 38F imaged in a single embryo over 6 h of development. Each data point represents a single nucleus over a 10-min time window, revealing the increase in the fraction of paired loci as development progresses. Data are separated into three plots for ease of visualization. (B) Nuclei from each timepoint were scored as “paired” if they fell within the shaded box in (A). Data were taken from three embryos each for transgenes at 38F (red) and 53F (blue) with error bars representing the standard error of the mean. For each button density *ρ*, we fitted the experimental pairing dynamics (Fig. 4 – Sup. Fig. 2A). Grey shading provides the envelope of the best predictions obtained for each *ρ* (dark grey) and its SD (light grey). (C) Phase diagram representing, as a function of *ρ*, the value of *E*_*p*_ (black line) that leads to the best fit between predicted and experimental developmental pairing dynamics. The predicted pairing strength is weaker than observed in the parameter space above the line, and stronger than observed below the line. Error bars represent the uncertainties on the value of *E*_*p*_ that minimizes the chi^2^-score at a given *ρ* value.

### 3) CONSTRAINING THE BUTTON MODEL USING DYNAMICAL MEASUREMENTS OF PAIRING PROBABILITY

Our button model predicts that the fraction of paired loci as a function of time depends on three parameters: the initial separation between homologous chromosomes *d*_*i*_, the density of buttons along the chromosome *ϱ*, and the button-button interaction energy *E*_*p*_ (Fig. 2D). As an initial test of our model, and to constrain the values of its parameters, we sought to compare model predictions to experimental measurements of the fraction of paired loci over developmental time.

Due to the still unknown molecular identity of the buttons, it was impossible to directly measure the button density and the button-button interaction energy. However, the initial separation between chromosomes *d*_*i*_ can be directly estimated using chromosome painting (Beliveau et al., 2012; Ried et al., 1998). To make this possible, we used Oligopaint probes (Beliveau et al., 2012) targeting chromosome arms 2L and 2R to perform chromosome painting on embryos ∼130 min after fertilization, corresponding to the beginning of cell cycle 14 (Foe, 1989) (Fig. 4 – Sup. Fig. 2A). The resulting distribution of distances between homologous chromosome territories was well described by a simple Gaussian distribution (Fig. 4 – Sup. Fig. 2B, red line).

We next investigated whether the button model quantitatively reproduced the pairing dynamics observed during development for reasonable values of the button density and the button-button interaction energy. We ran a series of simulations for various values of button density *ϱ* (from 10% to 100%) and strength of interaction *E*_*p*_ (from −0.5k_B_T to −5k_B_T) starting from values for the initial distance between homologous chromosomes *d*_*i*_ drawn from the inferred Gaussian distribution from our Oligopaint measurements (Fig. 4 – Sup. Fig. 2B, black line). For each parameter set, we monitored pairing dynamics as a function of developmental time and computed the average probability for a locus to be paired (Fig. 4 – Sup. Fig. 3A, black points) using the same criterion as in Figure 3E. By minimizing a chi^2^-score (Fig. 4 – Sup. Fig. 3B) between the predictions and the experimental pairing probability (Materials and Methods), we inferred, for each button density, the strength of interaction that best fits the data (Fig. 4C). Interestingly, the goodness of fit was mainly independent of button density (Fig. 4 – Sup. Fig. 3C): denser buttons require less strength of interaction to reach the same best fit (black line in Fig. 4C).

The inferred developmental dynamics quantitatively recapitulated the experimental observations for both investigated loci (Fig. 4B) for any choice of parameters given by the curve in Figure 4C. Note that our simulations did not predict the large increase in pairing observed for 38F between 5.5 h and 6 h (Fig. 4B). This disagreement may be a consequence of the proximity of 38F to the highly paired histone locus body (Fung et al., 1998; Hiraoka et al., 1993). In sum, the button model recapitulates the observed average pairing dynamics for a wide range of possible button densities coupled with interaction energies that are consistent with protein-DNA interactions.

### 4) PARAMETER-FREE PREDICTION OF INDIVIDUAL PAIRING DYNAMICS

The fit of our button model to the fraction of paired loci during development in living embryos (Fig. 4B) revealed a dependency between the interaction strength *E*_*p*_ and the button density *ϱ* (Fig. 4C). As a critical test of the model’s predictive power, we sought to go beyond averaged pairing dynamics and used the model to compute the pairing dynamics of individual loci. As can be seen qualitatively in the kymographs predicted by the model (Fig. 2B and Fig. 2 – Sup. Fig. 1), pairing spreads rapidly (within tens of minutes) from the buttons that constitute the initial points of contact along the chromosome. As a result, the button-model predicts that homologous loci undergo a rapid transition to the paired state as the zipping mechanism of pairing progression moves across the chromosome.

To quantify the predicted pairing dynamics of homologous loci, we collected single-locus traces containing individual pairing events from our simulations (Fig. 5A, top), which we defined as traces in which the inter-homolog distance drops below 0.65 µm for at least 4 min (Materials and Methods). For traces corresponding to each set of simulations with various values of *ϱ* and *E*_*p*_ (Fig. 5B), we calculated the median dynamics of inter-homolog distances around the pairing event. Across many values of *ϱ* and *E*_*p*_, the medians of the predicted trajectories leading up to the pairing event were very similar, with inter-homolog distances decreasing rapidly from 1-2 µm to below 0.65 µm at an accelerating rate over the course of 10-20 minutes (Fig. 5C-F). However, we do observe subtle differences in this pre-pairing stage: for a given button density, a stronger interaction energy *E*_*p*_ leads to a faster approach of the homologs (Fig.5 – Sup. Fig. 2). The nearly independence of this first period of the pairing dynamics with respect to *ϱ* and *E*_*p*_ suggests that the initial approach of homologous loci is mainly diffusion-limited, while there is a slight acceleration of pairing for stronger interactions energies due to an enhanced zippering effect.

**Figure 5.**
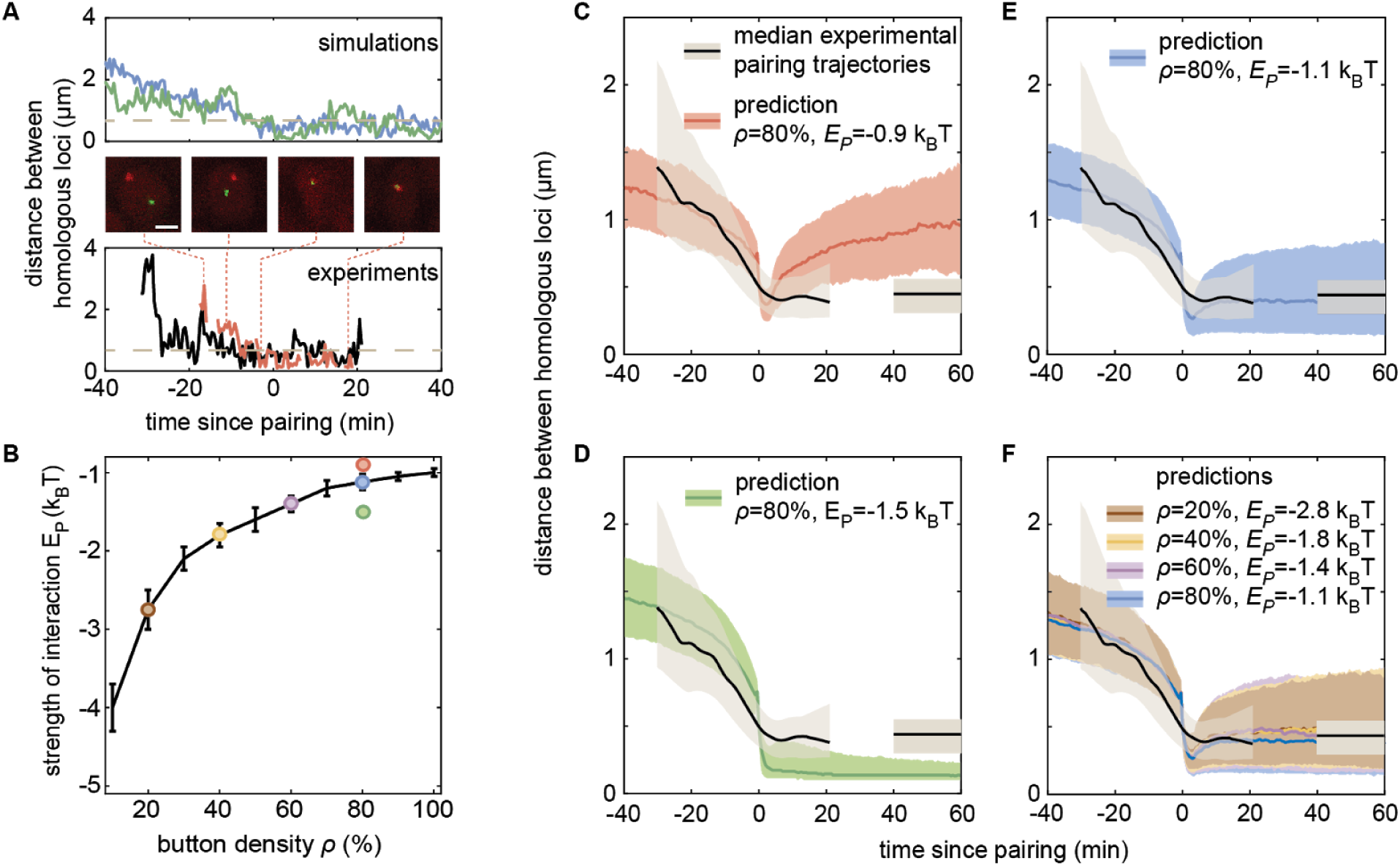
The homologous button model predicts individual pairing dynamics. (A) Examples of simulated (top) and experimental (bottom) pairing trajectories showing rapid transitions from the unpaired to the paired state. Simulations were carried out using *ϱ = 50%* and *E*_*p*_ = − 1.75 *k*_*B*_ *T*. See also Fig. 5 – Sup. Fig. 1. Scale bar is 2 μm. Dotted line in each graph represents our distance threshold of 0.65 µm used for aligning pairing traces. (B) Parameter range inferred from pairing probability dynamics in Figure 4 (black line), and parameters used for the simulations in C-F (color points). (C-F) Median pairing dynamics obtained from individual pairing trajectories detected during our experiments (black lines) and simulations (colored lines). Traces are centered at the time of pairing (time=0) defined as the time point where the inter-homolog distance decreases below 0.65 µm (grey dashed lines in (A)) for at least 4 min. The long-term, experimentally measured inter-homolog distance is plotted as a straight black line at the right of each panel. The first and third quartiles of the distribution of distances between homologous loci are indicated by the shaded regions (n=14 nuclei for experiments, n>10,000 traces for simulations).

In contrast to the initial pairing dynamics, varying model parameter values had a clear effect on the distance dynamics that followed the pairing event (Fig.5 – Sup. Fig. 2). Specifically, simulations with a weak *E*_*p*_ led to a slow increase in inter-homolog distances as time progressed (Fig. 5C, red), consistent with unstable pairing events. Conversely, simulations with a strong *E*_*p*_ were associated with tight pairing of homologous loci following the pairing event, with inter-homolog distances stably maintained around 130 nm, close to the spatial resolution of the model (Fig. 5D, green). Notably, the values of *ϱ* and *E*_*p*_ that best fit the averaged temporal evolution of the fraction of paired loci over development (Fig. 4 and 5B) all led to similar predictions for the median inter-homolog distance dynamics associated with pairing events. These traces converged to a stable long-term median inter-homolog distance of ∼0.5 µm (Fig. 5 E,F), which is nearly identical to the experimentally determined distance of ∼0.44 µm between homologous loci in stably paired nuclei (compare the colored and black lines in Fig. 5 E,F). Our results thus suggest that the slow dynamics of the pairing probability observed during development (Fig. 4) and the dynamics of inter-homolog distance after a pairing event (Fig. 5) are strongly correlated.

We then compared our simulated traces to experimental observations of pairing events in nuclei of live embryos. Among the movies that we monitored, we captured 14 pairing events matching the criteria of initial large inter-homolog distances that drop below 0.65 µm for at least 4 min (Fig. 5A; Fig. 5 – Sup. Fig. 1; Supplementary Movie 7). We aligned each of these pairing events using the same approach as with the simulated data described above, and calculated the median dynamics of inter-homolog distances around the pairing event (Fig. 5C-F, black lines). The pre-pairing dynamics were fully compatible with model predictions, with a rapid decrease in inter-homolog distances over 10-20 min (Fig. 5C-F). Further, the experimental post-pairing dynamics in inter-homolog distance were closely recapitulated (Fig. 5F) by the predictions made using parameters that best fit the pairing probability over developmental time (Fig. 5B). In sum, simulations of chromosomal behavior based on a button model with a defined set of parameters quantitatively recapitulate experimental observations of pairing events at individual loci, of stably paired homologs following a pairing event, and of the global progression of pairing dynamics over developmental time.

## Discussion

Since its discovery by Nettie Stevens over 100 years ago (Stevens, 1908), somatic homolog pairing has represented a fascinating puzzle for geneticists and cell biologists alike. The dissection of the molecular origins of somatic pairing presents a tractable case study to further our understanding of the 3D organization of chromosomes and the functional consequences of transactions among otherwise distant DNA loci. However, despite decades of research, the molecular mechanisms underlying somatic homolog pairing have remained elusive (Joyce et al., 2016). In this paper, we augmented the emerging button-based cartoon model of somatic homolog pairing by turning it into a precise theoretical model that makes quantitative and testable predictions of pairing dynamics as a function of the density of buttons throughout the chromosome and the specific interaction energy between buttons.

To assess the feasibility of this button model, we used it to predict chromosomal dynamics and then tested those predictions experimentally by tracking pairing dynamics at individual chromosomal loci in living embryos. Simulations predicted rapid transitions from unpaired to paired states resulting from a ‘zippering’ effect across the chromosomes where buttons that become paired via random encounters promote and stabilize pairing of adjacent buttons (Fig. 2B). The model predicts that this spread of pairing from button to button along the length of the chromosome ultimately leads to the formation of paired homologous chromosomes that remained stably associated throughout the rest of the simulation. Interestingly, this zippering effect driven by homologous buttons has also been suggested as a possible mechanism for meiotic pairing (Marshall and Fung, 2016).

In tracking pairing dynamics through early development in living embryos, we found quantitative agreement with the button model predictions: the transition from an unpaired to a paired state is a rapid event that occurs in just a few minutes (Fig. 5A), and paired chromosomal loci remain stably paired over the observation time of our experiments, up to 45 min (data not shown). Overall, the close quantitative agreement between observation and theory validates the button model as a mechanism that supports the initiation and maintenance of somatic homolog pairing. Further, our measurements constrain the range of possible values of the button density and interaction energy (Fig. 4C).

Two caveats may be considered in interpreting our analysis. First, our method of tracking homologous loci in living embryos relies on visualizing nascent RNAs generated from transgenes (Materials and Methods) rather than direct observation of DNA or DNA-binding proteins. While nascent RNAs provide a robust and convenient signal for the position of the underlying DNA (Chen et al., 2018; Lim et al., 2018), the method limits us to examining the behavior of transcriptionally active loci, which could behave differently from silent chromatin. In addition, our analysis could overestimate inter-homolog distances in paired nuclei if, for example, nascent RNA molecules from separate chromosomes are prevented from intermixing (Fay and Anderson, 2018). Second, our simulations do not account for complex behaviors of the genome that take place during development and that may also influence pairing dynamics and stability, including cell-cycle progression and mitosis (Foe, 1989), establishment of chromatin types and associated nuclear compartments (Hug et al., 2017; Ogiyama et al., 2018; Sexton et al., 2012; Yuan and O’Farrell, 2016), and additional nuclear organelles such as the histone locus body (Liu et al., 2006; White et al., 2011). Further testing and refinement of our theoretical and molecular understanding of somatic homolog pairing will require new approaches to incorporate the potential influences of these genomic behaviors in a developmental context.

A previous analysis of pairing and transvection in living embryos focused on the blastoderm phase, coinciding with the earliest developmental timepoints in our analysis, and found that inter-homolog interactions were generally unstable at that time (Lim et al., 2018). Thus, the embryo appears to transition from an early state that antagonizes stable pairing prior to cellularization to one that supports stable pairing at later time points of development. Prior studies have postulated changes in cell-cycle dynamics (Fung et al., 1998; Gemkow et al., 1998), chromatin states (Bateman and Wu, 2008a), or proteins that promote or antagonize pairing (Bateman et al., 2012a; Hartl et al., 2008; Joyce et al., 2012; Rowley et al., 2019) as potentially mediating a shift to stable pairing during the maternal-to-zygotic transition that occurs during blastoderm cellularization. Our data suggest that these changes mediate their effect on pairing by directly or effectively modulating button activity.

What is the molecular nature of the buttons? Prior studies based on Hi-C methods reported that the *Drosophila* genome contains distinct regions or peaks of tight pairing between homologs distributed with a typical density of 60-70% throughout the chromosome, which could represent pairing buttons (AlHaj Abed et al., 2019; Erceg et al., 2019; Rowley et al., 2019). Given such a button density and our experimental observations, our model predicts that a specific interaction energy between buttons would be ∼1-2 k_B_T (Fig. 4C), a value consistent with both typical protein-protein interactions (Phillips et al., 2012) and with electrostatic interactions between homologous DNA duplexes (Gladyshev and Kleckner, 2014; Kornyshev and Leikin, 2001). Moreover, two studies independently found enrichment for DNA-binding architectural and insulator proteins in tight-pairing regions (AlHaj Abed et al., 2019; Rowley et al., 2019), suggesting a potential role for these proteins in button function. In support of this view, an analysis of ectopically induced pairing *in vivo* found that chromosomal segments carrying candidate buttons are enriched in clusters of insulator proteins (Viets et al., 2019), and incorporation of insulator sequences into transgenes can stabilize pairing and transvection (Fujioka et al., 2016; Lim et al., 2018; Piwko et al., 2019). Notably, our analysis revealed a requirement for some degree of specificity between homologous buttons (Discussion – Sup. Fig. 1A,B), since simulations of non-specific interactions between buttons did not result in robust pairing (Fig. 2 – Sup. Fig. 2E). Perhaps a “code” of interactions between unique combinations of insulators and architectural proteins (Rowley et al., 2019; Viets et al., 2019) conveys the necessary specificity between homologous buttons for efficient pairing (Discussion – Sup. Fig. 1A,C).

While somatic homolog pairing is widespread in *Drosophila* and other Dipterans, it is curious that pairing of homologous sequences is rare in the somatic cells of other diploid species. It is possible that the sequences and proteins that underlie buttons are unique to Dipterans and are not present on the chromosomes of other species, perhaps due to the diversity of architectural proteins carried in the *Drosophila* genome (Cubeñas-Potts and Corces, 2015). Alternatively, chromosomes of other species may have the capacity to pair through encoded buttons, but are prevented from doing so by the functions of proteins that antagonize pairing, such as the condensin II complex (Hartl et al., 2008; Joyce et al., 2012; Rowley et al., 2019). However, most other diploid species do show a capacity to pair homologous chromosomes during the early stages of meiosis. While it is possible that meiotic pairing could be mediated via the same types of buttons that we postulate here (Marshall and Fung, 2016), our data do not directly address this question. Indeed, important differences appear to exist in the progression of meiotic pairing relative to somatic pairing, such as the highly dynamic and unstable associations between homologous loci (Ding et al., 2004) and rapid meiotic prophase chromosome movements (Lee et al., 2012) that have been observed in yeast, as well as unique chromosomal regions called pairing centers in *Caenorhabditis elegans* (MacQueen et al., 2005; Phillips et al., 2005). Therefore, multiple molecular mechanisms may accomplish the goal of aligning homologous chromosomes in different cellular contexts.

Importantly, our biophysical model of the otherwise cartoon-like button model coupled with quantitative live-cell imaging of pairing dynamics establishes a foundational framework for uncovering the parameters of button density and binding energy underlying somatic homolog pairing. In the future, we anticipate that our model will be instrumental in identifying and characterizing candidate button loci, and in determining how these parameters are modulated in the mutant backgrounds that affect pairing (Bateman et al., 2012a; Gemkow et al., 1998; Hartl et al., 2008; Joyce et al., 2012). Thus, our study significantly advances our understanding of the century-old mystery of somatic homolog pairing and provides a theory-guided path for uncovering its molecular underpinnings.

## Materials and Methods

### The homologous button model

We modeled two pairs of homologous chromosome arms as semi-flexible self-avoiding polymers. Each chromosome consists of *N*=3,200 beads, with each bead containing 10 kbp and being of size *b* nm. The four polymers moved on a face-centered-cubic lattice of size *L*_*x*_ *x L*_*y*_ *x L*_*z*_ under periodic boundary conditions to account for confinement by other chromosomes. Previously, we showed that TAD and compartment formation may be quantitatively explained by epigenetic-driven interactions between loci sharing the same local chromatin state (Ghosh and Jost, 2018; Jost et al., 2014). However, such weak interactions cannot lead to global homologous pairing (Pal et al. 2019). Here, to simplify our model, we neglect these types of interactions (whose effects are mainly at the TAD-scale) to focus on the effect of homolog-specific interactions. However, we do consider HP1-mediated interactions between (peri)centromeric regions that are thought to impact the global large-scale organization inside nuclei (Fig. 2 – Supp. Fig. 2D) (Strom et al., 2017).

Homologous pairing was modeled as contact interactions between some homologous monomers, the so-called buttons (Fig. 2A). For each pair of homologous chromosomes, positions along the genome were randomly selected as buttons with a probability *ϱ*. Each 10-kbp bead *i* of chromosome *chr* is therefore characterized by a state *p*_*chr,i*_ with *p*_*chr,i*_*=1* if it is a button (=0 otherwise) and *p*_*chr,i*_*= p*_*chr’,i*_ =1 if *chr* and *chr’* are homologous. In addition, the first 1,000 monomers of each chromosome were modeled as self-attracting centromeric and pericentromeric regions, the rest as neutral euchromatic regions. The energy of a given configuration was given by

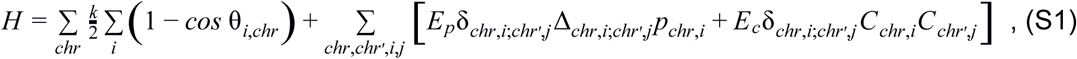

where *k* is the bending rigidity, *θ*_*i,chr*_ is the angle between the bond vectors *i* and *i+1* of chromosome *chr*, δ_*chr,i*;*chr*′,*j*_ = 1 if beads *i* from chromosome *chr* and *j* from *chr’* occupy nearest-neighbour sites (=0 otherwise), Δ_*chr,i*;*chr*′,*j*_ =1 if *i=j* and *chr* and *chr’* are homologous (=0 otherwise), *C*_*chr,i*_ =1 if bead *i* of *chr* is a (peri-)centromeric regions, *E*_*p*_ <0 is the contact energy between homologous buttons, *E*_*c*_ <0 is the contact energy between centromeric beads.

The dynamics of the chains followed a simple kinetic Monte-Carlo scheme with local moves using a Metropolis criterion applied to *H*. The values of *k* (*=1*.*5kT*), *b* (*=105* nm), *E*_*c*_ (*=−0*.*1kT*), *L*_*x*_*= L*_*y*_ (*=2* µm) and *L*_*z*_ (*=4* µm) were fixed using the coarse-graining and time-mapping strategies developed in (Ghosh and Jost, 2018) for a 10-nm fiber model and a volumic density=0.009 bp/nm^3^ typical of *Drosophila* nuclei. For every set of remaining parameters (the button density *ϱ* and the strength of pairing interaction *E*_*p*_), 250 independent trajectories were simulated starting from compact, knot-free, Rabl-like initial configurations (Dernburg et al., 1996): all centromeric regions were localized at random positions at the bottom of the simulation box, the rest of the chain being confined into a cylinder of diameter ∼600 nm and height ∼2 µm pointing toward the top of the box (see examples in Fig. 2C and Supplementary Movies 1, 2). The distance between the centers of mass of homologous chromosomes, noted as *d*_*i*_, typically varied between 0.5 µm and 3 µm. Each trajectory represented ∼10 h of real time. To model the developmental pairing dynamics, we ran simulations in which *d*_*i*_ was sampled from the distribution inferred from chromosome painting experiments (Figure 4 – Supplementary Figure 2).

To constrain model parameters, we compared the measured pairing dynamics (Fig. 4B) to the model prediction. Specifically, for each parameter set, we computed a chi^2^-score between the predicted dynamics and experimental timepoints

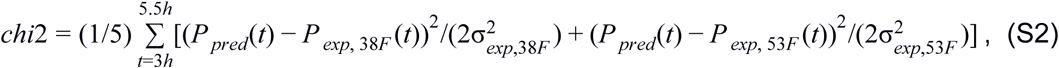

with *P* _*pred*_ (*t*) the predicted dynamics, *P*_*exp*, 38*F*_ (*t*) and *P*_*exp*, 53*F*_ (*t*) the experimental average dynamics for loci 38F and 53F, respectively, and σ_*exp*,38*F*_ and σ_*exp*,53*F*_ their corresponding standard deviations.

### DNA constructs and fly lines

Flies expressing a nuclear MCP-NLS-mCherry under the control of the nanos promoter were previously described (Bothma et al., 2018). To create flies expressing PCP-NoNLS-GFP, the plasmid pCASPER4-pNOS-eGFP-PCP-αTub3’UTR was constructed by replacing the MCP coding region of pCASPER4-pNOS-NoNLS-eGFP-MCP-αTub3’UTR (Garcia et al., 2013) with the coding region of PCP (Larson et al., 2011). Transgenic lines were established via standard P-element transgenesis (Spradling and Rubin, 1982). To create flies expressing MS2 or PP7 loops under the control of UAS, we started from plasmids piB-hbP2-P2P-lacZ-MS2-24x-αTub3’UTR (Garcia et al., 2013) and piB-hbP2-P2P-lacZ-PP7-24x-αTub3’UTR, the latter of which was created by replacing the MS2 sequence of the former with the PP7 stem loop sequence (Larson et al., 2011). The *hunchback* P2P promoter was removed from these plasmids and replaced by 10 copies of the UAS upstream activator sequences (Brand and Perrimon, 1993) and the *Drosophila* Synthetic Core Promoter (DSCP) (Pfeiffer et al., 2010). Recombinase-mediated cassette Exchange (Bateman et al., 2006) was then used to place each construct at two landing sites in polytene positions 38F and 53F (Bateman and Wu, 2008b; Bateman et al., 2012b). Flies carrying the GAL4 driver *nullo-GAL4*, which drives expression in all somatic cells during the cellular blastoderm stage of cell cycle 14, were a gift from Jason Palladino and Barbara Mellone. Flies carrying the GAL4 driver *R38A04-GAL4*, which drives expression in epidermal cells in germband-extended embryos (Jenett et al., 2012), were acquired from the Bloomington Drosophila Stock Center. Finally, the interlaced MS2 and PP7 loops under the control of the *hunchback* P2 enhancer and promoter (P2P-MS2/PP7-lacZ) were based on a previously described sequence (Wu et al., 2014).

To create embryos for analysis of pairing, mothers of genotype *10XUAS-DSCP-MS2; MCP-mCherry-NLS, PCP-GFP* were crossed to males of genotype *nullo-GAL4, 10XUAS-DSCP-PP7*. The resulting embryos are loaded with MCP-mCherry-NLS and PCP-GFP proteins due to maternal expression via the nanos promoter, and zygotic expression of *nullo-GAL4* drives transcription of MS2 and PP7 loops in all somatic cells starting approximately 30 min into cell cycle 14 (cellular blastoderm.) For pairing analysis, both MS2 and PP7 transgenes were in the same genomic location, either position 38F or 53F, whereas for a negative control, MS2 loops were located at 38F and PP7 loops were located at 53F. To visualize pairing at later times in development, the mothers indicated above were instead crossed to males of genotype *10XUAS-DSCP-PP7; R38A04-GAL4*, where both MS2 and PP7 loops were located at position 38F. Finally, to visualize MS2 and PP7 loops derived from the same genomic location, mothers of genotype *MCP-mCherry-NLS, PCP-GFP* were crossed to P2P-MS2/PP7-lacZ located at position 38F.

### Embryo preparation and image acquisition

Embryos were collected at 25 °C on apple juice plates and prepared for imaging as previously described (Garcia et al., 2013). Mounted embryos were imaged using a Leica SP8 confocal microscope, with fluorescence from mCherry and eGFP collected sequentially to minimize channel crosstalk. For each movie, the imaging window was 54.3 × 54.3 µm at a resolution of 768 × 768 pixels, with slices in each z-series separated by 0.4 µm. Z-stacks were collected through either 10 or 12 µm in the z plane (26 or 31 images per stack), resulting in a time resolution of approximately 27 or 31 s per stack using a scanning speed of 800 Hz and a bidirectional scan head with no averaging. For the pairing data in Figure 3, the imaging window was centered laterally for embryos that were pre-gastrulation; for post-gastrulation embryos, the imaging window centered on a dorsal view of the embryonic head region covering mitotic domains 18 and 20 (Foe, 1989), which shows minimal movements during gastrulation and germ band extension relative to other regions of the embryo. We compared pairing levels in these cells at 6 h of development to that of cells in a posterior abdominal segment at the same time point and found them to be nearly identical (75.0% paired, n=16 for anterior cells vs. 73.7%, n=19 in posterior cells according to the definition of pairing in Figure 3E), confirming that cells from different regions of the embryo are roughly equivalent for pairing dynamics at this stage. For positive control embryos with interlaced PP7 and MS2 loops driven by the *hunchback* promoter, embryos were imaged during cell cycle 13 and early cell cycle 14, and the imaging window was positioned laterally as previously described (Garcia et al., 2013). To assess pairing in late stage embryos using the *R38A04-GAL4* driver, embryos were aged to approximately 11-12 h and the imaging window was positioned laterally over an abdominal segment. For the developmental time course movies in Figure 4, imaging centered on mitotic domains 18 and 20 when these cells were in interphase. During timepoints when these domains were undergoing mitosis, an adjacent mitotic domain in interphase was imaged.

### Image analysis

All images were first run through the ImageJ plug-in Trainable Weka Segmentation (Arganda-Carreras et al., 2017) and filtered with custom classifiers to generate two separate channels of 3D segmented images that isolated fluorescent spots. These segmented spots were then fitted to a Gaussian with a nonlinear least squares regression to find the 2D center. Image z-stacks were then searched for any spots tracked for three or more contiguous z-slices and the rest were discarded. Additional manual curation was employed to confirm the accuracy of segmented images and to add any spots that were missed. An initial estimate of the center of each spot was set based on the z-slice in which the spot had the greatest maximum intensity within a predefined radius from its 2D center. These initial estimates were then used to seed a 3D Gaussian fit for each spot, the center of which was used for all distance calculation. This granted us not only sub-pixel resolution in x-y but also sub z-slice resolution, allowing for more precision in the z coordinate, which would otherwise be limited by the 0.4 µm spacing between consecutive stacked images created by confocal imaging.

Raw image z-stacks for each time frame were also maximum projected in the channel containing nuclearly localized MCP-mCherry to create 2D maps of all the nuclei in frame. These nuclear projections were then segmented and tracked in Matlab, followed by manual curation to ensure that each nucleus was consistently followed. One tracked particle lineage from each channel was then assigned a distinct nucleus based on its proximity to that nucleus in the 2D map and the particles in each channel were considered homologous chromosomes of one another. Since absolute coordinates of assigned particles were not possible to obtain due to cellular rotation and motion, all distance calculations were done with the relative coordinates of each locus from its homolog; any cellular rotation or motion was assumed to be conserved between loci in the same cell.

For the data in Figure 3, we qualitatively scored each nucleus based on the measured distances between red and green signals over the time that the signals were observed: “paired” nuclei showed small distances and little variation over time, and “unpaired” nuclei showed larger distances and greater variation over time. Nuclei that showed a transition from large distances and variation at earlier time points to smaller distances and variation at later time points were scored as “pairing” traces, and were not included in Figure 3 (see Figure 5). In assessing the stability of the paired state, we included both “paired” (n=25) and “pairing” (n=13) nuclei from three embryos in the total number of nuclei (n=38) assessed. In this analysis, we conservatively only included the observation time of “paired” nuclei (> 8 h of observation with no transition back to the unpaired state), although “pairing” nuclei also remained in the paired state throughout the remaining observation time once they became paired.

To align the traces presented in Figure 5 based on a timepoint when the loci become paired, we manually aligned all traces that had been qualitatively assessed as “pairing” traces according to several values of threshold distance and consecutive frames below that threshold. We then optimized this exploration for values that provided qualitatively good alignment of traces but that excluded as few traces as possible in order to maximize the data available for analysis. The same criteria were applied to identify and align pairing traces from simulations. All image analysis was done using custom scripts in Matlab 2019b unless otherwise stated.

### Chromosome painting

Embryos of genotype *w*^*1118*^ were aged to 2-3 h after embryo deposition, fixed, and subjected to DNA Fluorescence *In Situ* Hybridization (DNA-FISH) using 400 pmol of Oligopaint probes (Beliveau et al., 2012) targeting 2L and 2R (200 pmol of each probe; (Rosin et al., 2018)) as previously described (Bateman and Wu, 2008a). Hybridized embryos were mounted in Vectashield mounting medium with DAPI (Vector Laboratories), and three-dimensional images were collected using a Leica SP8 confocal microscope. To establish initial inter-homolog distances, an image from an embryo in early interphase 14 (as judged by nuclear elongation (Fung et al., 1998)) and with high signal-to-noise ratio was analyzed using the TANGO image analysis plugin for ImageJ (Belevich et al., 2016; Ollion et al., 2013, 2015). After segmentation and assignment of each painted territory to a parent nucleus, distances between territories were measured from centroid to centroid in 3D. Since homologous chromosomes are labeled with the same color, when territories produce a continuous region of fluorescence, a distance of zero was assigned. A total of 48 nuclei were analyzed for each of 2L and 2R.

## Acknowledgments

We thank Florian Jug for help with an earlier version of the nuclear tracking software. We also thank Francesco Ferrari, Gary Karpen, Cédric Vaillant, and members of the Jost group for fruitful discussions. HGG was supported by the Burroughs Wellcome Fund Career Award at the Scientific Interface, the Sloan Research Foundation, the Human Frontiers Science Program, the Searle Scholars Program, the Shurl & Kay Curci Foundation, the Hellman Foundation, the NIH Director’s New Innovator Award (DP2 OD024541-01), and an NSF CAREER Award (1652236). DJ acknowledges Agence Nationale pour la Recherche (ANR-18-CE12-0006-03, ANR-18-CE45-0022-01) and ITMO Cancer (Plan Cancer 2014-2019, Biologie des Systèmes n°BIO2015-08) for funding and CIMENT infrastructure (supported by the Rhone-Alpes region, Grant CPER07 13 CIRA) for computational resources. JRB was supported by grants from the National Institutes of Health (P20 GM0103423 and R15 GM132896-01), and an NSF CAREER Award (1349779).

## Supplementary Materials

### Supplementary Figures

**Figure 2 – Supplementary Figure 1.**
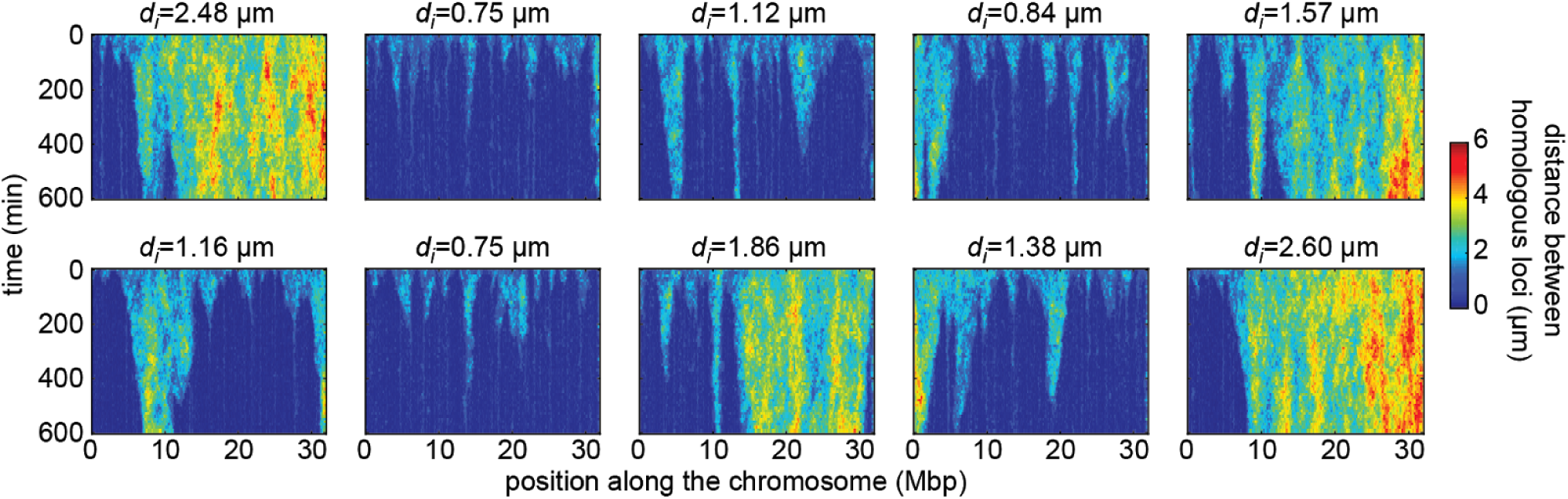
Simulated time evolution of distance between homologous loci. Examples of kymographs of the time-evolution of the distances between homologous regions predicted by the model in several simulated stochastic trajectories with different initial distances *d*_*i*_ for a button density of *ϱ=*65% and an interaction strength of *E*_*p*_ =−1.6k_B_T. Pairing operates via a “zipper” mechanism: as one pair of homologous loci becomes stably paired, the pairing of adjacent buttons is facilitated, leading to the “spreading” of pairing along the chromosome (flame-like patterns in the kymographs). For smaller initial distances, pairing is faster and more efficient.

**Figure 2 – Supplementary Figure 2.**
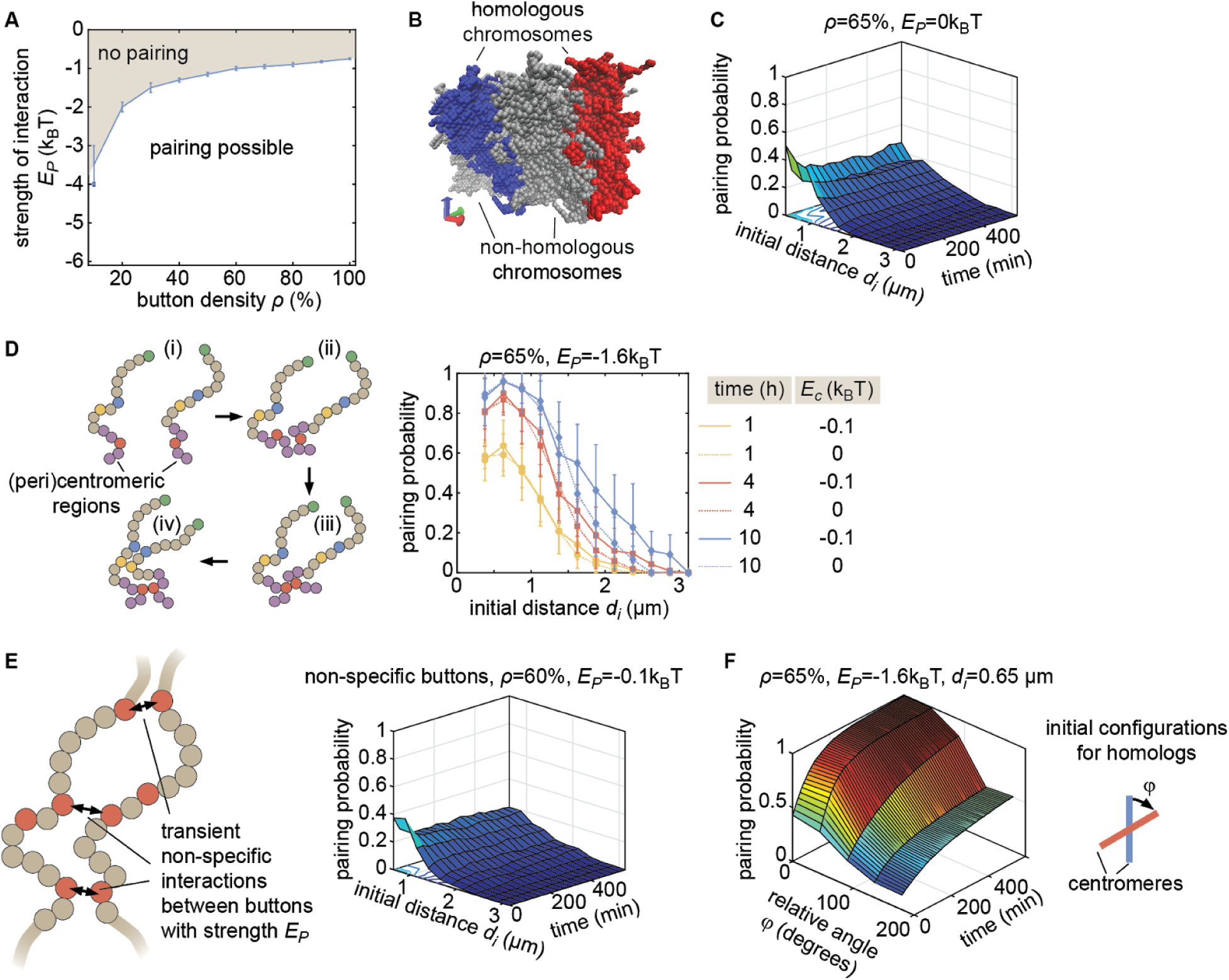
Properties of the button model. (A) Phase diagram of parameter ranges resulting in simulations compatible (white area) and incompatible (grey area) with pairing, starting from favorable initial homologous chromosome configurations (aligned chromosomes with *d*_*i*_ < 1 µm, as in Fig. 2C). (B) Representative simulation snapshot for *ρ*=65%and *E*_*p*_ =− 1.6 kT. When homologous chromosomes (red and blue monomers) are initially distant, steric hindrance by other chromosomes (grey monomers) may prevent pairing. (C) Behavior in the absence of specific interaction (*E*_*p*_ =0) in the homologous button model. Predicted average pairing probability between euchromatic homologous loci (considered as paired if distance ≤ 1μ*m*) as a function of time and the initial distance *d*_*i*_ between homologous chromosomes. (D) Impact of HP1-mediated interaction between (peri)centromeric regions. (Left) Pairing between chromosomes that are initially distant (i) is facilitated and accelerated by the non-specific interactions between centromeric regions that lead to the clustering (ii) of HP1 regions. This clustering facilitates the pairing of homologous buttons located in centromeric regions (iii) and the subsequent pairing via the zipping process of nearby euchromatic regions (iv). (Right) Predicted average pairing probability between euchromatic homologous loci (considered as paired if distance ≤ 1μ*m*) as a function of the initial distance *d*_*i*_ between homologous chromosomes at 1 h, 4 h, and 10 h for *ϱ=*65% and *E*_*p*_ =−1.6kT in the presence (*E*_*c*_ =− 0.1*kT*, full lines) or absence (*E*_*c*_ = 0*kT*, dashed lines) of interactions between (peri)centromeric regions. Even for large initial distances (*d*_*i*_ ≥ 2.5 μ*m*), we observe a weak but significant amount of pairing in the euchromatic regions. These predictions are consistent with DNA FISH experiments at various loci, suggesting an average higher pairing probability in heterochromatic loci during embryogenesis (see Fig.1 and Sup.Fig 2 in (Erceg et al., 2019)). (E) The non-specific button model. To verify whether a non-specific button model can lead to global pairing, we relaxed the homologous model to include specific attraction only between homologous loci and generated models where buttons may interact with any other buttons in the nucleus (left). In this model, the energy of a given configuration was described by

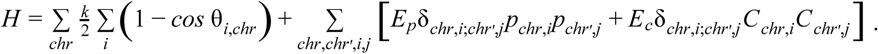

We varied *ϱ* from 10 to 100% and *E*_*p*_ from −0.025 to −4kT, one realization of which is shown here (right), without observing any global pairing of homologous chromosomes. Other parameters were as in the homologous button model (see Materials and Methods of the main text). Contacts preferentially form between buttons belonging to the same chromosome, or more weakly, between buttons of different chromosomes but not necessarily between homologous loci. (F) Effect of the relative initial orientation between homologs. Predicted average pairing probability between euchromatic homologous loci (considered as paired if distance ≤ 1μ*m*) as a function of time and the angle φ between the initial configurations of homologs (see inset), for an initial distance between the centers of mass of homologous chromosomes *d*_*i*_ = 0.65 μ*m*. We see that lower relative angles lead to more efficient pairing, with Rabl-like configurations corresponding to 0° ≤ φ ≤ 45°.

**Figure 3 – Supplementary Figure 1.**
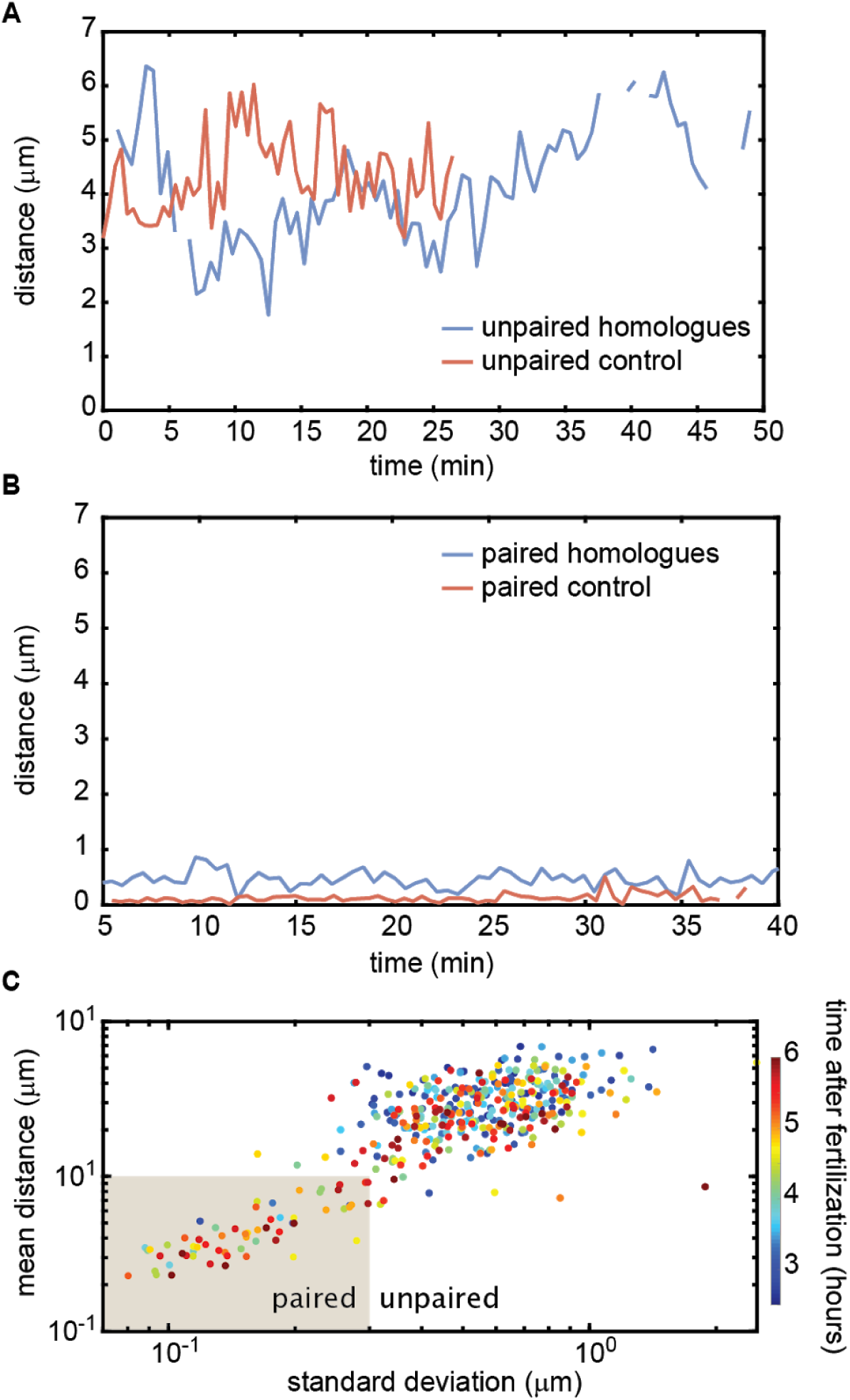
Homologous chromosome reporters inserted at the 53F genomic location. (A) Representative traces of the dynamics of the distance between imaged loci for unpaired homologous loci and the negative control. (B) Representative traces of the dynamics of the distance between imaged loci for paired homologous loci and the positive control. The traces in (A) and (B) are comparable to those measured at the 38F genomic location shown in Figure 3C,D. (C) Mean vs. standard deviation of the separation of each pair of homologous loci imaged in a single embryo over 6 h of development (analyses of 3 embryos are combined). Each data point represents a 10-min window.

**Figure 4 – Supplementary Figure 1.**
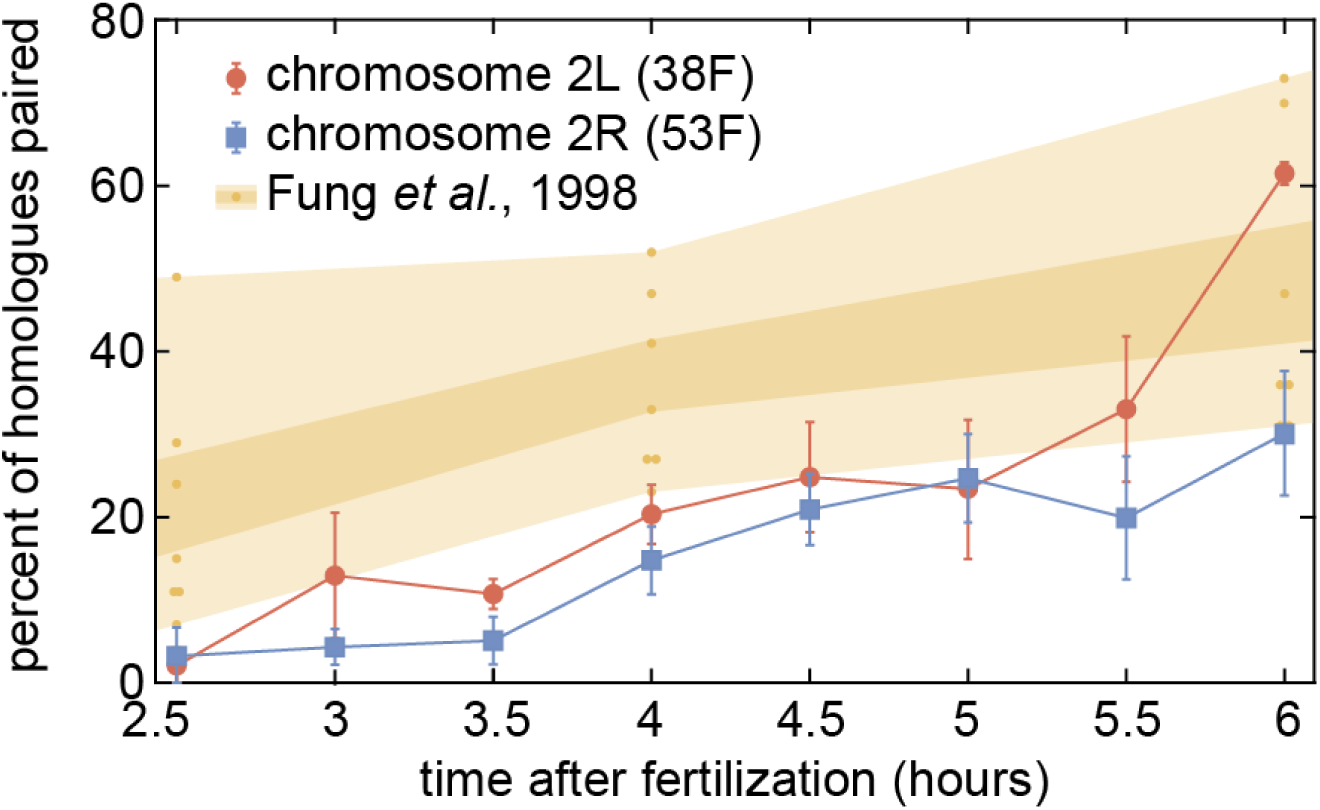
Comparison of our pairing data to previous results. Pairing dynamics measured by live-imaging at two chromosomal loci (red and blue points) as presented in Figure 4B. The progression of pairing observed in fixed embryos using DNA-FISH was obtained from (Fung et al., 1998); orange points represent the pairing measured at seven euchromatic loci on chromosome arm 2L at three timepoints, and orange shading represents standard error (dark) and range (light). Our data is consistent with, but systematically lower than the range of pairing observed in the prior study.

**Figure 4 – Supplementary Figure 2.**
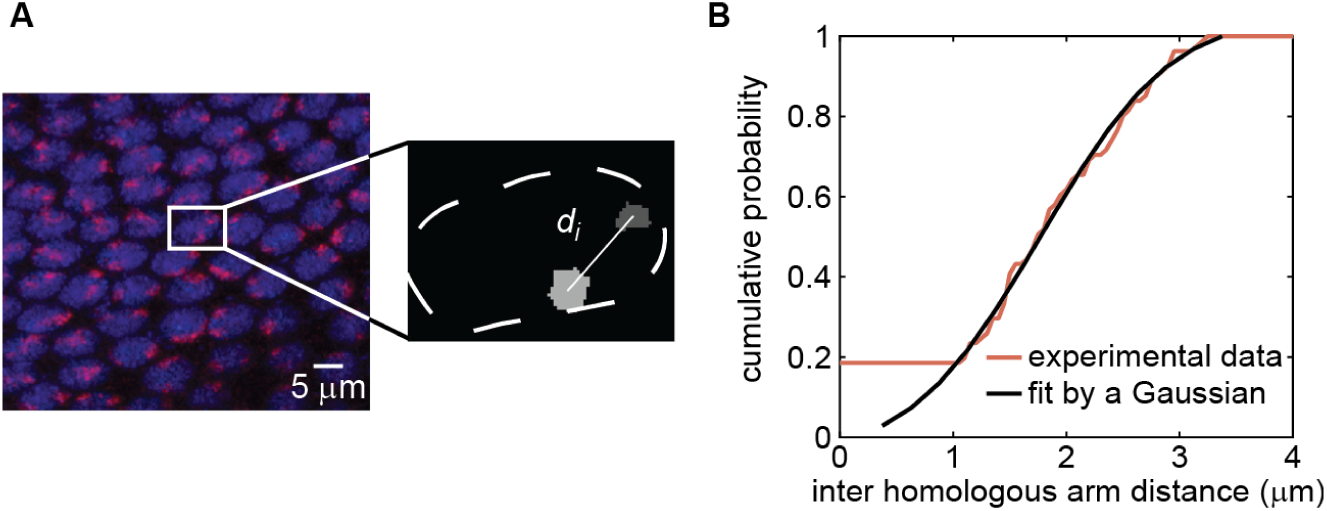
Establishing values for initial distance *d*_*i*_ between homologous chromosomes via chromosome painting. (A) Chromosome painting of chromosome arm 2L (red), carrying transgene location 38F, in an embryo in early cycle 14. Nuclei (blue) are stained with DAPI. Inter-homolog distances were determined by segmenting painted regions in 3D using ImageJ and measuring center-to-center distances (inset; see Materials and Methods for details). Chromosome arm 2R, carrying transgene location 53F, was sampled in the same field of cells using a different fluorescent tag (not shown). (B) Distribution of inter-homologous arm distances measured from 48 nuclei in the image in (A) (red curve). Measurements from chromosome arms 2L and 2R were completely overlapping and therefore were combined. The curve is nicely fitted by a Gaussian distribution (black curve, mean=1.9, SD=0.9).

**Figure 4 – Supplementary Figure 3.**
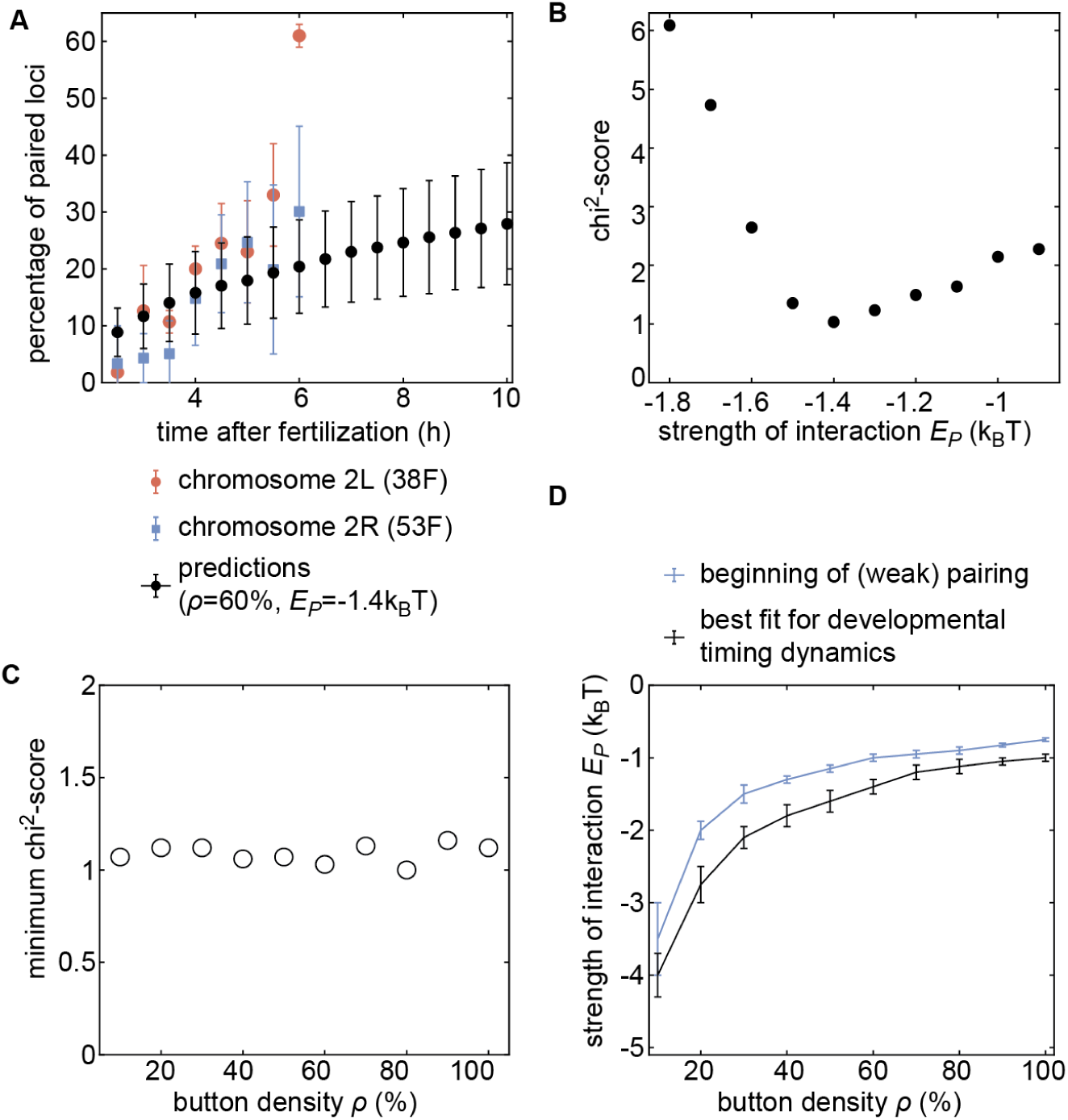
Inference of developmental pairing dynamics. (A) Predicted time evolution of the mean pairing dynamics computed over all buttons in the simulation (black dots) for a distribution of initial inter-homolog distances *d*_*i*_ given in Fig. 4A and for *ϱ*=60% and *E*_*p*_ =−1.4 k_B_T. Error bars in simulated data represent SD of the pairing dynamics computed over all the buttons (n = 3,200), while error bars in experimental data represent the standard error of the mean (n = 3 embryos for each chromosomal position). The model can recapitulate the experimentally measured pairing dynamics. (B) Evolution of the chi^2^-score as a function of *E*_*p*_ for *ϱ*=60%. For some *E*_*p*_ values, we performed two sets of independent simulations. (C) The values of the minimum chi^2^-score obtained for all the investigated button densities are very similar, suggesting an equivalent predicting power for every *ϱ*. (D) Phase diagram representing, for each button density *ρ*, the value of *E*_*p*_ (black line) that leads to the best fit between predicted and experimental developmental pairing dynamics. The blue line stands for the weak pairing transition defined in Figure 2 – Supplementary Figure 2A.

**Figure 5 – Supplementary Figure 1.**
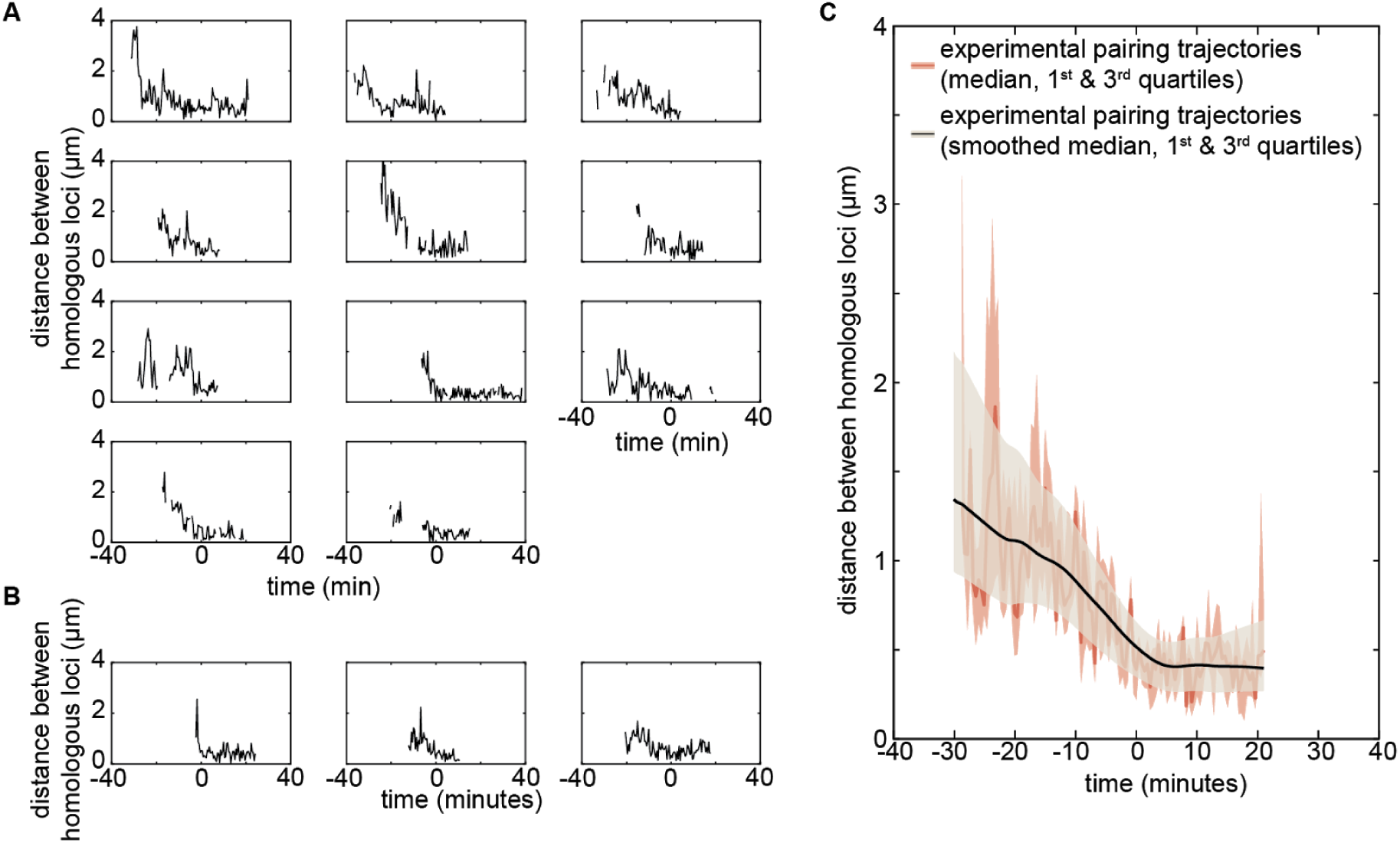
Experimental individual pairing dynamics. (A,B) Fast single-locus pairing dynamics illustrated by individual pairing traces detected for the experiments on locus 38F (A, n=11) and locus 53F (B, n=3) centered at the time of pairing (time=0) defined as the time point where the inter-homolog distance decreases below 0.65 µm for at least 4 min. (C) From the 14 individual pairing trajectories observed experimentally, we estimated at each time point the median (dark red full line) and first and third quartiles (light red dashed lines) of the distribution of distances between homologous loci. The corresponding smoothed median is plotted, and the 1st and 3rd quartiles (as in Figure 5 of the main text) are plotted in black and grey shaded areas. Smoothing was performed using the *smooth* function of Matlab (method=*lowess*, span=*40*).

**Figure 5 – Supplementary Figure 2.**
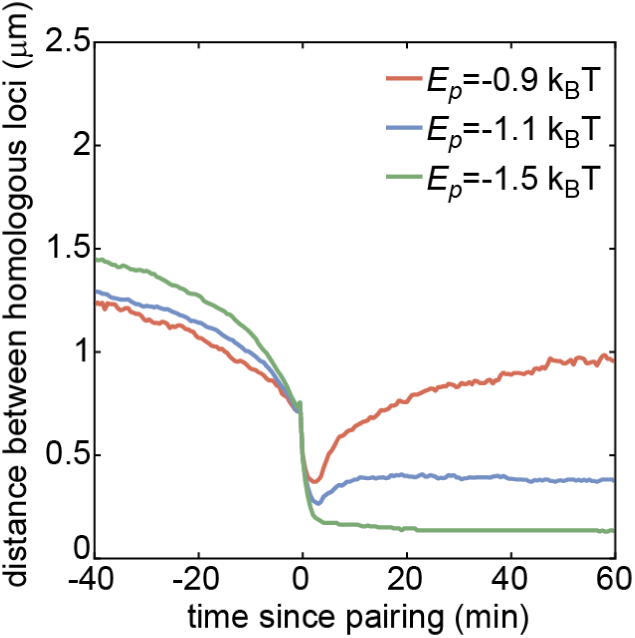
Median pairing dynamics obtained from individual pairing trajectories detected during our simulations (colored lines) for *ϱ*=80% and different values of *E*_*p*_. Traces are centered at the time of pairing (time=0) defined as the time point where the inter-homolog distance decreases below 0.65 µm for at least 4 min. While the main differences lie in the post-pairing period, with stronger *E*_*p*_ leading to a smaller distance between paired homologous loci, weak changes in the pre-pairing dynamics can be observed with stronger *E*_*p*_ leading to faster decay in the inter-homolog distance.

**Discussion – Supplementary Figure 1.**
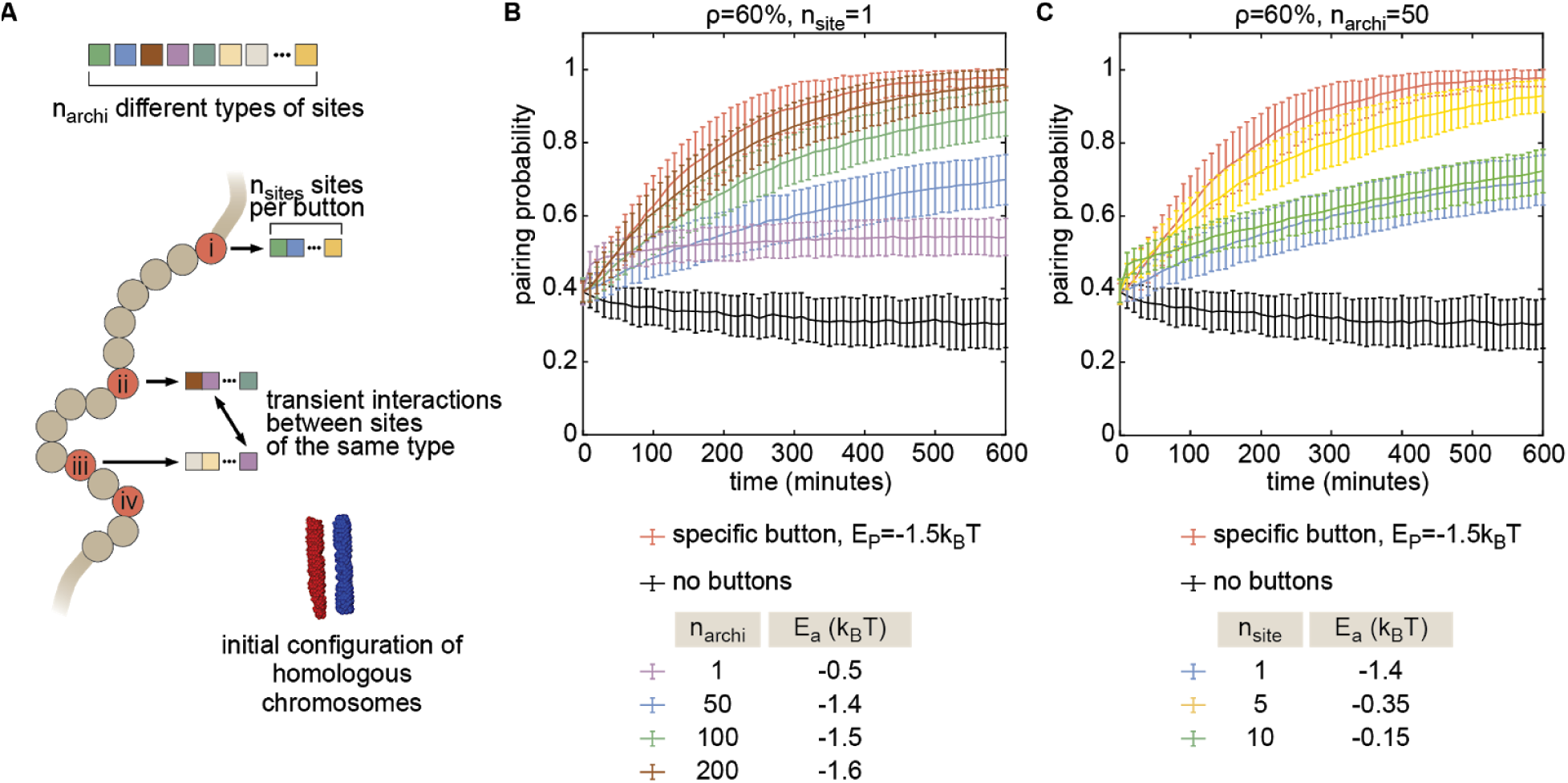
The combinatorial button model. (A) To investigate the possible nature of buttons, we developed a more general model in which buttons are made of *n*_*site*_ binding sites for specific architectural proteins. Every site is occupied by one type of architectural proteins randomly chosen among *n*_*archi*_ types. Two buttons may interact if they have common binding sites for some architectural proteins. The energy of a configuration is given by 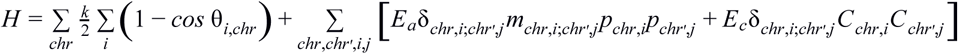, with parameters defined as in Equation S1 and *m*_*chr,i*;*chr*′,*j*_ the number of common binding sites between buttons *(chr,i)* and *(chr’,j)*, and *E*_*a*_ < 0 the strength of interaction between binding sites bound to the same architectural proteins. Homologs share the same pattern of binding sites, i.e., *m*_*chr,i*;*chr*′,*i*_ = *n*_*site*_ if *chr* and *chr’* are homologous and if *i* is a button. For example, the case *n*_*site*_=1, *n*_*archi*_=1 (*m*_*chr,i*;*chr*′,*j*_ =1, *E*_*a*_ = *E*_*p*_) corresponds to the non-specific button model described in Fig. 4 – Sup.Fig. 2E. To simplify and to avoid potential issues arising from periodic boundary conditions, we focused on favorable situations for pairing where homologs are initially aligned and close to each other (∼640 nm between their respective centers of mass) (see inset in A) and where the monomers evolve in a close box (rigid wall conditions). Using this model, we first investigated how specific a button should be in order to lead to pairing. We fixed *n*_*site*_=1 and varied *n*_*archi*_ and *E*_*a*_ for a button density of 60%. In these cases, *n*_*archi*_ represents the number of button types. In (B), we plotted, for each *n*_*archi*_, the time evolution of the average pairing probability between homologous sites (paired if distance ≤ 1μ*m*) for the *E*_*a*_-value that leads to maximal pairing. As a point of comparison, we also plotted the corresponding curves in the absence of buttons (black line) and for the homologous button model investigated in the main text (red line) for the *E*_*p*_-value (−1.5kT) consistent with experiments at the corresponding button density. We observed that as *n*_*archi*_ increases (as the buttons become more specific) the pairing efficiency increases. The full specific model (red) becomes well approximated by our combinatorial button model for *n*_*archi*_>200, i.e., when the number of buttons for one type is less than 10 per chromosome at 10kbp resolution. This suggests that pairing needs a significant degree of specificity via a large number of button types but each button type may be present in a small amount. However, we consider it unlikely that there are enough different architectural proteins in *Drosophila* to reach such single-site specificity. A possibility to increase specificity from a small number of proteins is to allow more than one binding site per button (*n*_*site*_>1). The number of different buttons is then (*n*_*site*_)!/[(*n*_*archi*_-*n*_*site*_)!(*n*_*site*_)!]. In (C), we fixed *n*_*archi*_=50, an upper maximal number of architectural proteins in flies, and varied *n*_*site*_. For each *n*_*site*_, we plotted the pairing probability for the *E*_*a*_-value that leads to maximal pairing. We observed that there exists an optimal number of sites (here ∼5) for which the pairing is close to the one obtained with the homologous button model. This corresponds to a value that leads to a large diversity of buttons while maintaining a low number of spurious interactions between non-homologous buttons, which is of the order of *n*_*site*_/*n*_*archi*_∼0.1.

## Supplementary Movies

**Supplementary Movie 1**.

Polymer simulations of homologous pairing. Example of a 4-h numerical simulation of the homologous button model (*ϱ*=60%, *E*_*p*_ =−1.6 k_B_T) with frames taken every 30 s. The movie focuses on one pair of homologs (red and blue polymers). Orange and cyan parts of these chains represent their (peri)centromeric regions. Surrounding transparent light-grey chains represent the periodic boundary images of the simulated chains. The scale bar is 1 µm. Homologous loci in closed contact (distance <200 nm) are colored in green. Initially, chromosomes are randomly placed in a Rabl-like configuration. Then, they decompact and dynamically evolve in a crowded environment. First pairing events are diffusion-driven, followed by a spreading of pairing to nearest buttons via a zippering effect.

**Supplementary Movie 2**.

Polymer simulations of homologous pairing. Same as in Supplementary Movie 1 but taken from a different simulation run. The two pairs of homologs are highlighted (red/blue for one pair; purple/dark blue). Orange/cyan and pink/light blue parts of these chains represent their (peri)centromeric regions. Surrounding transparent light-grey chains represent the periodic boundary images of the simulated chains. The scale bar is 1 µm. Homologous loci in closed contact (distance <200 nm) are colored in green.

**Supplementary Movie 3**.

Representative confocal movie of a live *Drosophila* embryo (cell cycle 14 to gastrulation) in which MS2 and PP7 loops are integrated at equivalent positions on homologous chromosomes. Examples of nuclei are highlighted whose loci display characteristic dynamics, including loci that do not pair (“Unpaired”), loci that are already paired (“Paired”), and loci that are observed transitioning from the unpaired to the paired state (“Pairing”). Image stacks were taken roughly every 30 s and max-projected for 2D viewing.

**Supplementary Movie 4**.

Representative confocal movie of a live *Drosophila* embryo (roughly 4.5 h old) in which MS2 and PP7 loops are integrated at equivalent positions on homologous chromosomes. A greater proportion of nuclei show paired homologs relative to the earlier time point represented in Supplementary Movie 3. Image stacks were taken roughly every 30 s and max-projected for 2D viewing.

**Supplementary Movie 5**.

Representative confocal movie of a live *Drosophila* embryo (roughly 5.5 h old) in which MS2 and PP7 loops are integrated at various positions on homologous chromosomes (MS2 at position 38F and PP7 at position 53F) where we expect no pairing between transgenes. Image stacks were taken roughly every 30 s and max-projected for 2D viewing.

**Supplementary Movie 6**.

Representative confocal movie of a live *Drosophila* embryo (cell cycle 14) in which MS2 and PP7 loops were interlaced in a single transgene on one chromosome at polytene position 38F to act as a positive control for pairing. Both GFP and mCherry are co-localized to the same locus in all transcriptional loci. Image stacks were taken roughly every 30 s and max-projected for 2D viewing.

**Supplementary Movie** 7.

Representative distance trajectory of two loci denoted as “pairing” (plotted in the black experimental trace in Figure 5A) showing a rapid transition from large distances at earlier time points to smaller distances at later time points. The trajectory is shown alongside movies of the nucleus and gastrulating embryo from which the distances were calculated to help visualize the speed at which this pairing occurs. Image stacks were taken roughly every 30 s and max-projected for 2D viewing.

